# Mapping distribution of invasive plant species and uncertainty using citizen science, remote sensing, and deep learning

**DOI:** 10.64898/2026.06.10.731341

**Authors:** Xu Qiang, Lauren E. Gillespie, Jia Xi, Dimitrios Gounaridis, Kai Zhu

## Abstract

Invasive plants pose a major environmental problem, threatening biodiversity, altering ecosystem functions, and causing economic loss. Climate change is altering environmental conditions, potentially facilitating the spread of invasive plant species, posing challenges for ecosystem management and biodiversity conservation. Accurate predictions of invasive species distributions are therefore essential for effective monitoring and early intervention. Species distribution models (SDMs) have become an important tool for predicting species habitats, but many studies rely on traditional machine learning approaches, focus on single-species predictions and overlook uncertainty associated with future climate scenarios. This study aims to evaluate the performance of a deep learning-based SDM framework, Deepbiosphere, for predicting both native and invasive plant species distributions on a regional scale, the US state of Michigan, and to assess how climate scenario uncertainty influences spatial predictions of invasive species risk particularly on two focal invasive species. Results show that Deepbiosphere outcompeted other baseline models by on average of 10.98% with a mean AUC-ROC of 0.79 across 1553 vascular plant species. For two invasive species *Rhamnus cathartica* and *Ailanthus altissima*, Deepbiosphere respectively improved modeling accuracy by an average of 56.41% and 74.99%, suggesting its enhanced predictive capability for invasive species. Current predictions indicated that *R. cathartica* is already broadly suitable across much of Michigan, whereas *A. altissima* is currently more restricted to southern regions. Under future climate scenarios, both species were projected to expand northward, with a particularly strong expansion signal for *A. altissima*. Prediction uncertainty was spatially heterogeneous, where general circulation models (GCMs) were the dominant source of uncertainty across most of the state. By integrating citizen science, remote sensing, and deep learning, we produced high-resolution risk-uncertainty maps for key invasive species and highlighted the importance of explicitly mapping uncertainty to support more informed invasive species management under climate change.

## 1 Introduction

Global biodiversity is facing unprecedented challenges due to climate change, habitat loss, invasive species, and other anthropogenic pressures (IPBES, 2019). Among these factors, invasive species are a major concern because they disrupt biodiversity by outcompeting native species through aggressive resource acquisition and fast growth (Dukes et al., 2009). The impact of invasive plants spans from ecological to societal and economic aspects. Ecological impacts include the loss of critical habitat for native species, food web interruption, alteration of soil chemistry and hydrological cycles, and microclimate shifts (Ehrenfield, 2001; Vilà et al., 2011).

Beyond these ecological impacts, invasive plants can also impact human health, damage recreational resources, and have led to massive economic loss of up to 120 billion dollars in the United States (Fantle-Lepczyk et al., 2022). Climate change, as another major driver affecting biodiversity, has complex interactive effects on invasive species. Under climate change, distributions of most invasive species are expected to expand due to their physiological advantage, phenological flexibility, and new invasion pathways, and such expansion of invasive species in return can reduce carbon sequestration and increases greenhouse gas emission, leading to a positive feedback loop (Clements & Ditommaso, 2011; Finch et al., 2021; Hellmann et al., 2008; Merow et al., 2017).

Managing invasive species under a changing climate presents various challenges that complicate prevention, control, and restoration strategies. For instance, elevated carbon dioxide levels, drought, and high temperatures have been shown to increase the tolerance of certain invasive plants to herbicides by hindering uptake and translocation of such chemical treatments (Manea et al., 2011). Furthermore, climate change can impede biological control by causing phenological mismatch between invasive plants and their insect predators or pathogens (Hellmann et al., 2008). Mechanical control, including cutting, girdling, and tilling, may also become more expensive and less effective if warmer winters lead to higher survival rates for invasive species. Even proactive strategies, such as assisted migration, may unintentionally introduce other species, creating a new dispersal pathway for invasive species (Finch et al., 2021). Additionally, invasive species are known to have high dispersal rates, rapid growth rates, and high tolerance to extreme environmental conditions, all of which exceed land managers’ capacity for timely detection and monitoring using traditional approaches (Cohen et al., 2024; Finch et al., 2021). Ultimately, these interacting drivers pose significant challenges to invasive species management. A first step in effective management is to accurately predict species distributions and how they will change under climate change. As demonstrated by Leung et al. (2002), the cost-benefit ratio of early detection and intervention far exceeds that of long-term management once an invasion has stabilized. In this context, by presenting spatially explicit predictions, species distribution models (SDMs) can play a fundamental role in early-stage monitoring to detect invasions quickly and thus guide actions to prevent or deter establishment even under the inherent uncertainties of global environmental change (Srivastava et al., 2021).

SDMs, also known as bioclimatic envelope models, ecological niche models, and habitat suitability models, are one of the most widely used methods in conservation science and ecological modeling (Elith et al., 2006; Franklin, 2010; Guisan et al., 2013). SDMs have been extensively studied for invasive plant species (Srivastava et al., 2019) at different geographical scales. This includes from identifying worldwide invasion hotspots that are suitable for a great number of invasive plants (Omer et al., 2026) and predicting an invasive Asteraceae shrub at the global scale (Mainali et el., 2015), to exploring the expansion of an invasive annual grass in the western U.S.(Nietupski et al., 2024) and two important woody invasive plants in East Africa (Eckert et al., 2020) at a regional scale.

SDMs typically use statistical and machine learning methods to produce empirical descriptions and spatial predictions of species distribution (Franklin, 2023; Valavi et al., 2022), by exploring the relationship between geographical occurrences of species and corresponding environmental variables (Guisan & Zimmermann, 2000). Before the development of deep learning (DL), these conventional non-DL models were often constrained in expressivity and scalability due to the complicated relationship between species and environment (Kellenberger et al., 2026). Many non-DL models rely on simple assumptions that might be insufficient to represent the nonlinear and complex relationships affected by biotic and abiotic interactions (Wisz et al., 2013). They frequently overlook spatial context, which means that predictor values are only extracted for the center observations, thereby ignoring the surrounding landscape structure (Lembrechts et al., 2019). As the amount and types of ecological datasets are increasingly growing, these non-DL models become less capable of leveraging and handling input data, thereby hindering their application on a broader scale (Kellenberger et al., 2026).

The advent of DL in the last few decades has brought the potential to address the above limitations. Compared to non-DL models, DL models have a greater ability to learn complex relationships as the dataset size and complexity increase. Some DL models can naturally take spatial context into account and compute or recombine the input predictors, which provides more information to predict the probability of presence. The modular design in DL architecture enables the integration of different input types and the prediction of multiple species simultaneously with common base neutral network layers (Kellenberger et al., 2026). Early studies mainly relied on feed-forward artificial neural networks (ANN), which demonstrated the potential of neural-network approaches for modeling species distributions (Benkendorf & Hawkins, 2020; Harris, 2015; Li & Wang, 2013; Zhang & Li, 2017; Zhang et al., 2020), but the architecture remained relatively simple, with only a few layers. However, more recent studies have delved deeper into the convolutional neural network (CNN) for SDMs. For example, Botella et al. (2018) used relatively advanced layers to explore CNN. Deneu et al. (2021) employed Inception V3, a successful CNN architecture for computer vision (Szegedy et al., 2016). Among these advances in DL-based SDMs, Deepbiosphere stands out because it combines a TResNet head (Ridnik et al., 2021) with an MLP head (Battey et al., 2020) to utilize remote sensing data and environmental variables, respectively, and has achieved extremely accurate performance in modeling plants in California, USA (Gillespie et al., 2024).

Despite the wide use of SDMs in ecology and conservation and the progression of DL-based SDMs, several important gaps remain. First, large-scale multispecies DL-based SDMs have rarely been applied to invasive plant species, even though invasive species management often requires broad geographic coverage and the ability to evaluate multiple species simultaneously. As investigated by Dutra Silva et al. (2021), from 1996 to 2019, there were a total of 526 articles addressing invasive species with SDMs involved. The majority of them still used non-DL single species approaches, with MaxEnt and Random Forest (RF) being the most frequently used. Although a few recent studies introduced joint SDMs or multispecies SDMs (JSDM or MSDM) (Ovaskainen & Abrego, 2020; Pollock et al., 2014; Sharma et al., 2025; Tikhonov et al., 2020), they still relied on a single framework built on non-DL approaches. To our knowledge, only two studies developed multispecies SDMs that were DL-based (Brun et al.,

2024; Gillespie et al., 2024), but this method has not been applied to invasive species analyses. Second, although SDMs are widely used for future projections under climate change, the outcomes merely focus on distribution maps or range shift (Poudel et al., 2025; Thapa et al., 2018), and uncertainty is less often analyzed and spatially presented (Thomas et al., 2024). This limits the ability to distinguish areas of varying levels of risk and uncertainty, which is important to inform management decision-making.

Building on the research gaps outlined above, this study addresses two research questions: 1) How effectively can a DL-based multispecies species distribution model map invasive plant distributions at the regional scale with high spatial resolution? 2) How does climate scenario uncertainty influence the spatial predictions of invasion risk? To answer these questions, this study compared Deepbiosphere with other baseline models to generate high-resolution statewide habitat suitability maps for two invasive plant species in the US state of Michigan under current and future climates. Ensemble mean suitability, prediction uncertainty, and risk–uncertainty classification were then used to identify areas of robust and uncertain invasion risk with potential relevance for management prioritization. An additive model to decompose variance was used to investigate the impact of climate scenarios on invasion risk.

## 2 Methods

### 2.1 Study area and target invasive species

Our study area focused on the land area of the state of Michigan, USA. Michigan has a humid temperate climate strongly moderated by the Great Lakes (Albert, 1995), which shaped its regional ecological pattern. Michigan is largely forested across the state but contains diverse ecosystems between the Upper Peninsula and the Lower Peninsula, with the southern Lower Peninsula dominated by deciduous forests and agricultural land, while mixed forests are more prevalent across the northern Lower Peninsula and Upper Peninsula. Invasive plant species occur statewide, with the highest concentrations observed in the Lower Peninsula (Cohen et al., 2024).

We chose two representative invasive species in Michigan, common buckthorn (*Rhamnus cathartica*) and tree of heaven (*Ailanthus altissima*), as the target species because of their severe ecological impacts on local plant communities. Common buckthorn is an invasive shrub native to Europe and Asia. It dominates disturbed and undisturbed areas, such as roadsides, pastures, old fields, and woodlots, and can be easily spread by birds carrying its seeds. It alters soil chemistry, promotes invasive earthworms, and acts as a critical host for major agricultural pests (Henghan et al., 2026.; MDAR, 2025; MISP, n.d.). Tree of heaven is a fast-growing and stress-tolerant invasive tree that outcompetes Michigan’s native plants through allelopathic toxins and aggressive root cloning. It often thrives in disturbed and urban environments, thus damaging infrastructure, and serves as the primary host for the spotted lanternfly, a pest that severely threatens Michigan’s agriculture (MISP, n.d.; Sladonja et al., 2015).

### 2.2 Data collection and preparation

We collected plant occurrence data across the Kingdom Plantae in Michigan from the Global Biodiversity Information Facility (GBIF.org, 2025), covering the period from 2015 to 2025. We filtered the records to include only those observed by humans with a coordinate uncertainty radius of less than or equal to 120 meters and no geospatially flagged issues. In total, 305,343 observations across 2705 species,1009 genera, and 252 families were downloaded. We filtered the dataset to include only the observations of vascular plants. Then, following the preprocessing steps of Gillespie et al. (2024), we conducted spatial thinning to reduce sampling bias by removing duplicate records of the same species within a 150 m radius and eliminating species whose observations were confined to a single 256 m radius. To create the joint occurrence dataset, we applied neighbor imputation, which added any additional species observations observed within an overlapping 256 m radius to a given observation (**Figure S1**). Finally, any species that contained fewer than 200 total observations was removed. This resulted in a dataset of 248498 total observations encompassing 1553 unique vascular plant species across 676 genera and 157 families.

For remote sensing data, we used imagery from the USDA’s National Agricultural Imagery Program (NAIP) acquired in 2012 for the entire state of Michigan (USGS Earth Resources Observation and Science Center, 2018). To train the CNN model, the images were partitioned into 256m×256m tiles, where each pixel represented a 1m×1m (**Figure S1**). The raw NAIP imageries in Michigan are projected into two projection coordinate systems, depending on their location. Therefore, we transformed all tiles in UTM Zone 16N into UTM Zone 17N. Then, we linked each species observation by its geographic coordinates to a 256m×256m tile with the observation centered in the image.

For the current climate, we downloaded the 19 bioclimatic variables from WorldClim Version 2 at 30 arc-second resolution, which is approximately 1km spatial resolution (Fick & Hijmans, 2017). All bioclimatic variables were standardized using z-score normalization to a mean of 0 and a standard deviation of 1 across Michigan.

For future climate projections, we obtained bioclimatic variables from multiple general circulation models (GCMs) under different Shared Socioeconomic Pathways (SSPs) for the mid-21st century (2041–2060). Future bioclimatic variables were also downloaded from WorldClim Version 2 at 30 arc-second resolution (Fick & Hijmans, 2017). To capture climate model uncertainty, we selected five GCMs that covered low to high climate sensitivity projections. The selected GCMs were MIROC6, BCC-CSM2-MR, MRI-ESM2-0, CMCC-ESM2, and IPSL-CM6A-LR. To capture emissions pathway uncertainty, we selected three SSPs representing low, intermediate, and high greenhouse gas emission trajectories: SSP1-2.6, SSP2-4.5, and SSP5-8.5. Variation in change in predicted habitat suitability across these combinations was interpreted as climate scenario uncertainty.

The administrative boundary of Michigan was obtained from the Global Administrative Areas database Version 4.1 (GADM) (2025), which would be used to define the study extent and clip all bioclimatic layers. Land cover data were derived from the National Land Cover Database (NLCD) (USGS, 2025), which classifies land cover across the United States at 30 m spatial resolution. The NLCD dataset was used to mask open water areas and retain only the land boundary within Michigan that would be used for prediction and visualization.

### 2.3 Dataset partition

As widely discussed, it is more likely that spatially close data points share similar attribute values because of the spatial dependency (Tobler, 1970; Hiimans, 2012). In the case of SDMs, geographically close data points often share similar environmental conditions and species compositions. To address this problem, we used two types of train-test splitting: uniform split and spatial cross-validation splits. The uniform split was to randomly select a subset of observations uniformly across Michigan as the test set, while the remaining observations were used for training. To mitigate data leakage due to spatial autocorrelation, we put all overlapping observations in the test set, so that the images and observation data of the test set were excluded from the train set. Besides, as the diameter of a pixel in bioclimatic raster could be up to 1200m converted from arc-second, we removed the observations that were located within 1300 m of observations in the test set, thus avoiding the same input value of bioclimatic variables in both the train and test sets.

In the absence of an independent validation dataset, we further used spatial cross-validation to test the model’s extrapolation ability and cross-validate the uniform split. The spatial cross-validation split involved a 7-fold cross-validation (**Figure S2**). The Michigan study region was partitioned into several latitudinal bands. In each fold of the spatial cross-validation, one 1-degree band was held out as the test region while the remaining bands were used for training (**Table S1**). As discussed above, training observations within 1300 m of the test region were also removed to prevent data leakage. Due to differences in observation density across geographic bands, the number of training and testing samples varied among 7 cross-validation folds.

### 2.4 Model training and evaluation

#### 2.4.1 Multimodal deep learning-Deepbiosphere

Deepbiosphere is a multi-species joint deep learning model that integrates citizen science observations with high-resolution imagery. The model consists of two main components designed to process different types of inputs and a multispecies structure to utilize co-occurrence labels. The TResNet head processes remote sensing data, while the MLP head processes bioclimatic variables (Gillespie et al., 2024). Overall, Deepbiosphere combines remote sensing data, species occurrence data containing taxonomic information and species co-occurrence patterns, and bioclimatic data to predict thousands of species simultaneously.

We trained all the Deepbiosphere models using a learning rate of 1×10 ^5^, 12 epochs, and a batch size of 150. We used a novel loss function, sampling-aware binary cross-entropy (BCE), to measure the discrepancy between predicted probabilities and observed species occurrence labels (Eq.1). Accounting for sampling bias in citizen science data, sampling-aware BCE has been demonstrated to perform well when dealing with multi-label image classification tasks because it ensures that present and absent classes contribute equally to the total loss. (Gillespie et al., 2024).

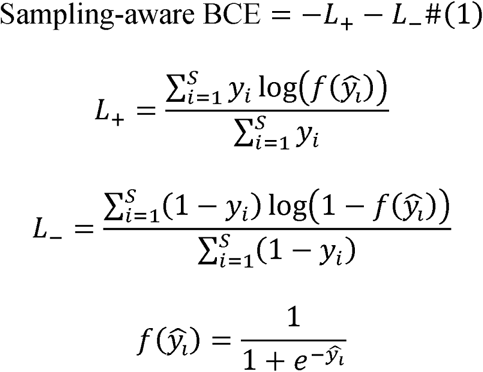

where *L*_+_and *L*_-_represent the normalized loss contributions from observed presences and absences (or background points), respectively.*y_i_*ɛ{0,1} denotes the ground truth label for species *i*, with *y_i_*=1.1indicating presence and *y_i_*=0 indicating absence or pseudo-absence.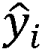 is the raw model output (logit) for species *i*, and 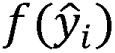 represents the predicted probability obtained using the sigmoid function. *S* is the total number of species. 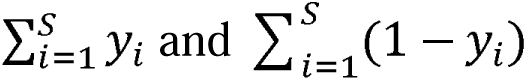 represent the number of present and absent species in a sample, respectively.

#### 2.4.2 Convolutional neural network-TResNet

To train a CNN-based SDM with NAIP imagery as the only input, we chose the modified TResNet architecture as another baseline model, which would be referred to as the RS TResNet. TResNet is a small CNN-based residual neural network, which is optimized for efficient GPU inference and is widely used for multi-label image classification tasks in computer vision (Ridnik et al., 2021). To incorporate the RGB and Near-Infared bands in NAIP imagery, the TResNet architecture was adapted to have 4 input channels.

We trained all RS TRestNet models with the same hyperparameters and loss as discussed above in the Deepbiosphere model. Training was conducted for 12 epochs with a batch size of 150 and a static learning rate of 1×10□□. The sampling-aware BCE loss function described above was also used for this model.

#### 2.4.3 Multilayered perceptron (MLP)

A multilayer perceptron model (MLP) was used as a baseline model to model nonlinear relationships between environmental predictors and species occurrence. This model, in our case, referred to as Bioclim MLP, was adapted from the architecture of Battey et al. (2020). The network comprises two fully connected layers with 1,000 neurons each, a dropout layer with a dropout rate of 0.25, and two additional layers with 2,000 neurons each (Gillespie et al., 2024).

We trained MLP models using the same hyperparameters and loss as the Deepbiosphere models for consistency. Training was conducted for 12 epochs with a batch size of 150 and a static learning rate of 1×10□□. The sampling-aware BCE loss function described above was also used for this model.

#### 2.4.4 Machine learning baselines

For comparison with Deepbiosphere, we employed the two most widely used machine learning SDMs, Random Forest (RF) (Breiman, 2001) and MaxEnt (Phillips et al., 2006). For each species, a separate model was trained one by one with only bioclimatic variables as input. Overall, 50,000 background points were generated using a circular sampling strategy centered on training observations. The radius of each sampling circle was defined as the median distance between occurrences of the focal species. For both the RF and MaxEnt models, an equal number of presence and background points was used to avoid class imbalance. For the RF model, 1,000 trees were produced using balanced bootstrap sampling. The MaxEnt model was implemented using the “no threshold” option to produce continuous probability values. All other model parameters were set to their default value.

#### 2.4.5 Performance evaluation

We utilized a wide variety of accuracy metrics calculated on predictions of the test set to evaluate the model’s performance. These accuracy metrics are divided into three broad categories: binary classification metrics, discrimination metrics, and ranking metrics. Binary classification metrics measure the model’s ability to correctly classify species presence or absence at a probability threshold of 0.5 in this study. Precision, recall, and F1 Score were calculated using per-species aggregation, while accuracy was referred to as presence accuracy and defined as the proportion of samples for which the species observed at a given location was correctly predicted as present. For each sample, only the single observed species associated with that location was evaluated. Presence accuracy was calculated following Gillespie et al. (2024)

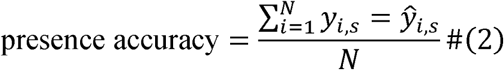

where *N* is the number of observations, *y*_i,s_is the ground truth presence (1) or absence (0) of species *s* at observation *i*, and 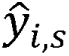 is the predicted binary presence or absence for species *s* at observation *i* after applying a threshold (≥ 0.5). The expression 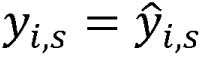 evaluates to 1 when the correct species observed at location *i* is predicted as present, and 0 as absent.

Discrimination metrics measure the model’s ability to distinguish between species presence and absence without an arbitrary threshold. Specifically, they are the Area Under the Receiver Operating Characteristic curve (AUC-ROC) and the Area Under the Precision-Recall Curve (AUC-PRC). Instead of setting a fixed threshold of 0.5, these metrics integrate model performance across a continuous probability threshold and were calculated by per-species aggregation. Ranking metrics focus on the ranking of a species by predicted probability for a given image or observation. Similarly, Top-100 accuracy, a single-label ranking metric, was calculated by per-species aggregation. To further assess the model’s ability to capture co-occurrence patterns, we also utilized a multi-label ranking metric called mean average precision (mAP). Except for presence accuracy, all other metrics were calculated using scikit-learn (Pedregosa et al., 2011).

Model evaluation used an early stopping strategy based on performance. The epoch of evaluation was selected as the epoch that achieved the highest mean AUC-ROC. After evaluation, the best-performing model and its corresponding optimal epoch were then used to generate species probability predictions for both the current climate and future climate scenarios.

### 2.5 Future climate scenario uncertainty

To evaluate the uncertainty associated with future climate projections, species distribution predictions of two target invasive species were generated under multiple climate scenarios by the best model. For each pixel, the ensemble mean and standard deviation (SD) of predicted probabilities were calculated across all scenarios, where the mean represents the central tendency and SD represents the variability among predictions of different scenarios (Eq. 3&4).

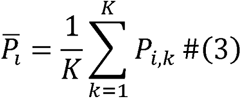

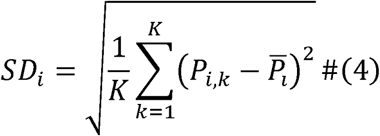

where 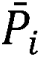 is the ensemble mean predicted probability at pixel *i,SD_i_* is the standard deviation of predicted probabilities at pixel *i,P_i,k_* is the predicted probability at pixel *i* under climate scenario *k*, and *Κ* is the total number of climate scenarios.

The ensemble mean probability or habitat suitability served as a proxy for predicted potential invasion risk. Pixels with mean probabilities ≥ 0.5 were classified as high-risk areas, whereas pixels with mean probabilities < 0.5 were classified as low-risk areas. To further visualize potential northward range expansion, we delineated the upper leading edge of predicted habitat suitability following the longitudinal band method of Zhu et al. (2012). For each vertical raster column, we identified the northernmost pixel with predicted probability ≥ 0.5. These pixels were then connected to represent the upper leading edge of suitable habitat.

Prediction uncertainty was quantified using the standard deviation of predicted probabilities across scenarios. SD values were categorized into three uncertainty levels based on the distribution of SD values. SD < 25th percentile was classified as low uncertainty, SD between the 25th and 75th percentiles was classified as moderate uncertainty, and SD > 75th percentile was classified as high uncertainty. Finally, invasion risk and prediction uncertainty were combined to construct a risk–uncertainty classification map containing six categories. This map provides a spatial representation of areas with different levels of predicted invasion risk and climate scenario uncertainty.

To quantify the change in habitat suitability and inform management, we defined the change (ΔP) as the difference between the ensemble mean of future suitability and current suitability (Eq.5).

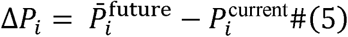

where *ΔP_i_*, is the change in predicted habitat suitability at pixel 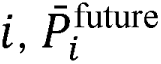is the ensemble mean predicted probability under future climate scenarios at pixel i,and *P_i_^current^* is the predicted probability under current climate conditions at pixel *i.*.

Pixels with ΔP< −0.1 were classified as decreasing, pixels with −0.1≤ΔP≤0.1were classified as stable, and pixels with ΔP>0.1were classified as increasing (Thomas et al., 2024). This classification was used to generate change maps and to summarize the spatial proportions of each change category for each species.

To assess the relative importance of different uncertainty sources habitat suitability change, for each pixel, scenario-based changes were first calculated for each GCM–SSP combination (Eq.6).

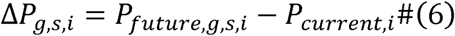

where Δ*P_g,s,i_* is the scenario-based change in predicted habitat suitability at pixel *i,P_future,g,s,i_* is the predicted future suitability at pixel *i* under GCM *g* and SSP *s*, and *Pcurrent,i* is the predicted current suitability at pixel *i*.

Then, we used a two-way ANOVA without an interaction term, which is also known as an additive model, to decompose the total variance, represented by the sum of squares, among scenario-based predictions into GCMs, SSPs, and residual components (Eq.7). Here, the ANOVA framework was used for variance decomposition rather than for statistical testing. The proportional contribution of each source was calculated relative to the total sum of squares (Eq.8). The dominant uncertainty source was defined as the component with the largest proportional contribution at each pixel.

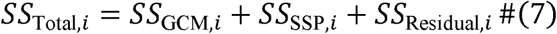

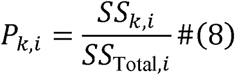

where *SS*_Total,*i*_ is the total sum of squares at pixel *i,SS_GCM,i_* is the sum of squares explained by differences among GCMs, *_SSP,i_* is the sum of squares explained by differences among SSPs, and *SS_Residual,i_* is the residual sum of squares.*P*_k,i_ denotes the proportional contribution of the uncertainty source *k* at pixel *i*, where *k* refers to GCM, SSP, or Residual.

## 3 Results

### 3.1 Overall model performance

Across all accuracy metrics evaluated on the uniform split, except for presence accuracy, Deepbiosphere consistently achieved the highest performance compared to both deep learning and traditional machine learning baseline models (**Table 1**; **Figure 2**). For the binary classification metrics, Deepbiosphere achieved the highest precision (0.0256), Recall (0.4999), and F1 score (0.0465), whose improvements ranged from 15.8% to 53.3%, from 0.1% to 60.9%, and from 18.3% to 51.5%, respectively. Although Deepbiosphere did not outcompete other baseline models on presence accuracy, its performance was still comparable to two DL-based models, around 0.83, and was much higher than two traditional machine learning models.

**Figure 1.**
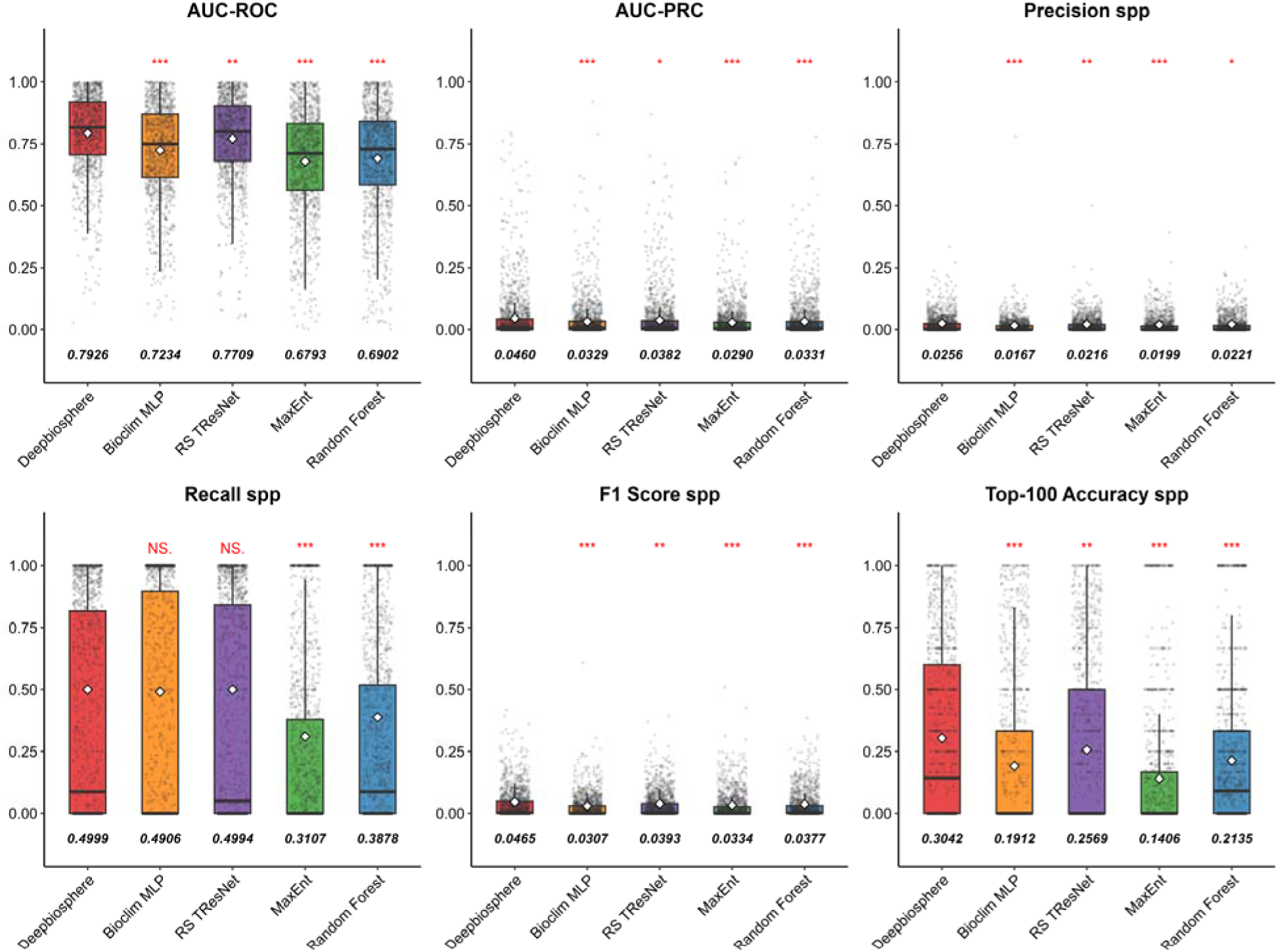
Comparison of the per-species accuracy metrics across shared 1123 species on the uniform split with the statistical results. White diamonds represent mean values that are annotated below each boxplot. Stars indicate results from an unpaired Student’s t-test, with *** indicating p-value <0.001, ** indicating p-value<0.01, * indicating a p-value <0.05, and NS. indicating a nonsignificant P-value > 0.1.

**Figure 2.**
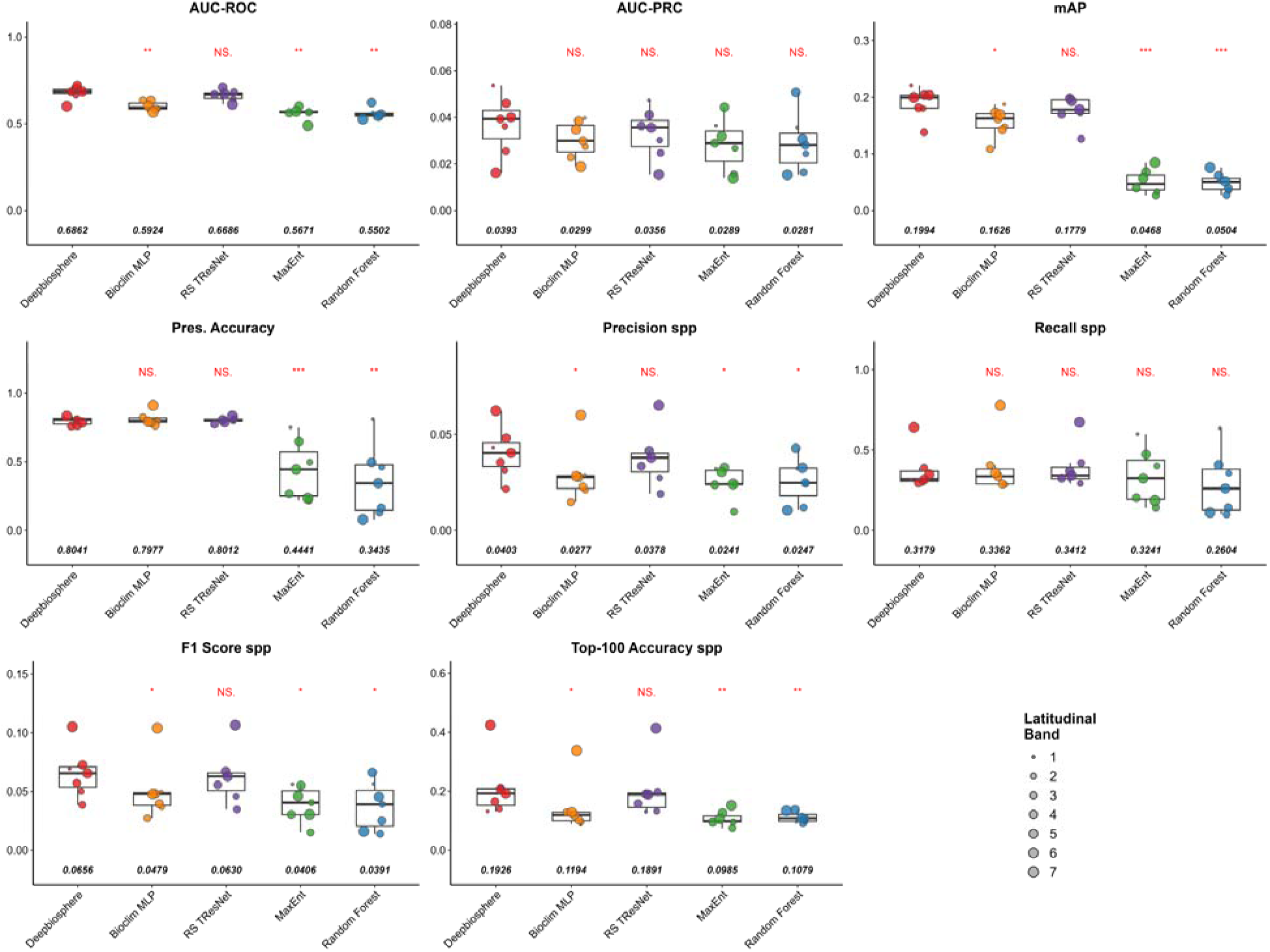
Comparison of accuracy metrics across 7 spatial cross-validation splits with statistical results. Median values are annotated below each boxplot. Stars indicate results from a Wilcoxon rank-sum test, with *** indicating p-value <0.001, ** indicating p-value<0.01, * indicating a p-value <0.05, and NS. indicating a nonsignificant P-value > 0.1.

**Table 1.** Comparison of accuracy metrics for species on the uniform split.

| Model name | AUC<br>ROC | AUC<br>PRC | mAP | Pre.<br>Accuracy | Precision<br>spp | Recall<sub>spp</sub> | F1<sub>spp</sub> | Top-100<sub>spp</sub> |
| --- | --- | --- | --- | --- | --- | --- | --- | --- |
| Deepbiosphere | **0.7926** | **0.0460** | **0.1459** | 0.8223 | **0.0256** | **0.4999** | **0.0465** | **0.3042** |
| BioClim MLP | 0.7234 | 0.0329 | 0.1102 | **0.8338** | 0.0167 | 0.4906 | 0.0307 | 0.1912 |
| RS TResNet | 0.7709 | 0.0382 | 0.139 | 0.832 | 0.0216 | 0.4994 | 0.0393 | 0.2569 |
| MaxEnt | 0.6793 | 0.029 | 0.0268 | 0.282 | 0.0199 | 0.3107 | 0.0334 | 0.1406 |
| Random Forest | 0.6902 | 0.0331 | 0.0348 | 0.3238 | 0.0221 | 0.3878 | 0.0377 | 0.2135 |

Means are reported for each metric, and bold entries are the highest value for each metric. Spp refers to the per-species level.

Deepbiosphere’s presence accuracy value was 0.8223, while BioClim MLP achieved the highest with a value of 0.8338, followed by RS TResNet, which achieved a value of 0.832. However, deep learning models performed better in terms of presence accuracy than machine learning models did. In terms of the most important discrimination metric, Deepbiosphere showed the highest AUC-ROC (0.7926) and AUC-PRC (0.0460), with improvements of mean AUC-ROC from 2.8% to 16.7%, with an average of 10.98%, indicating a largely improved ability to distinguish species presences across a gradient of thresholds.

Improvements were most pronounced in ranking metrics. Deepbiosphere also achieved the highest Top-100 accuracy (0.3042) and the highest mAP (0.1459). It improved Top-100 accuracy by 18.4% to 116.4% and improved mAP by 5% to 444.4%, compared with baseline models. Boxplots of per-species metrics further support these findings, showing higher mean values for Deepbiosphere across metrics (**Figure 2**). Statistical significance was assessed across species using t-tests, showing that Deepbiosphere significantly outperformed baseline models for most metrics (p < 0.05) across species, including AUC-ROC, AUC-PRC, precision, F1 score, and Top-100 accuracy (**Figure 2**).

To test the model’s extrapolation capacity and predict species distribution in unknown areas, we conducted model evaluation across spatial cross-validation bands. Due to the extremely imbalanced number of samples and distribution shift in Band 7’s training and testing set (**Figure S3&S4**), Band 7 exhibited outlier behavior relative to other spatial bands. Therefore, we reported medians to avoid the influence of extreme values. Similar to the evaluation on uniform split, Deepbiosphere also consistently achieved the strongest performance across most accuracy metrics, except for Recall, where RS TResNet had the best performance (**Table 2**). In terms of discrimination ability, Deepbiosphere achieved the highest median AUC-ROC (0.6862), significantly exceeding BioClim MLP (0.5924, p < 0.01), MaxEnt (0.5671, p < 0.01), and RF (0.5502, p < 0.01), while performance was comparable to RS TResNet (0.6686, p = NS.), suggesting that remote sensing imagery alone captures much of the discriminative signal. For binary metrics, Deepbiosphere achieved the highest median presence accuracy (0.8041), Precision (0.0403), and F1 (0.0656), with significant differences against most baselines, though Recall showed no significant differences across models. Noticing that, the difference in each binary metric between Deepbiosphere and RS TResNet was tested as not significant, indicating the comparable performance of these two models. Deepbiosphere significantly performed better than other models, except for RS TResNet, on presence accuracy, precision, and F1 (**Figure 3**). A similar pattern occurred in ranking metrics. Deepbiosphere achieved the highest mAP (0.1994) and Top-100 accuracy (0.1926), and its mAP and Top-100 accuracy were significantly higher than those of BioClim MLP, MaxEnt, and RF (p < 0.05), while there was no significant difference in both ranking metrics between Deepbiosphere and RS TResNet (**Table 2**; **Figure 3**).

**Figure 3.**
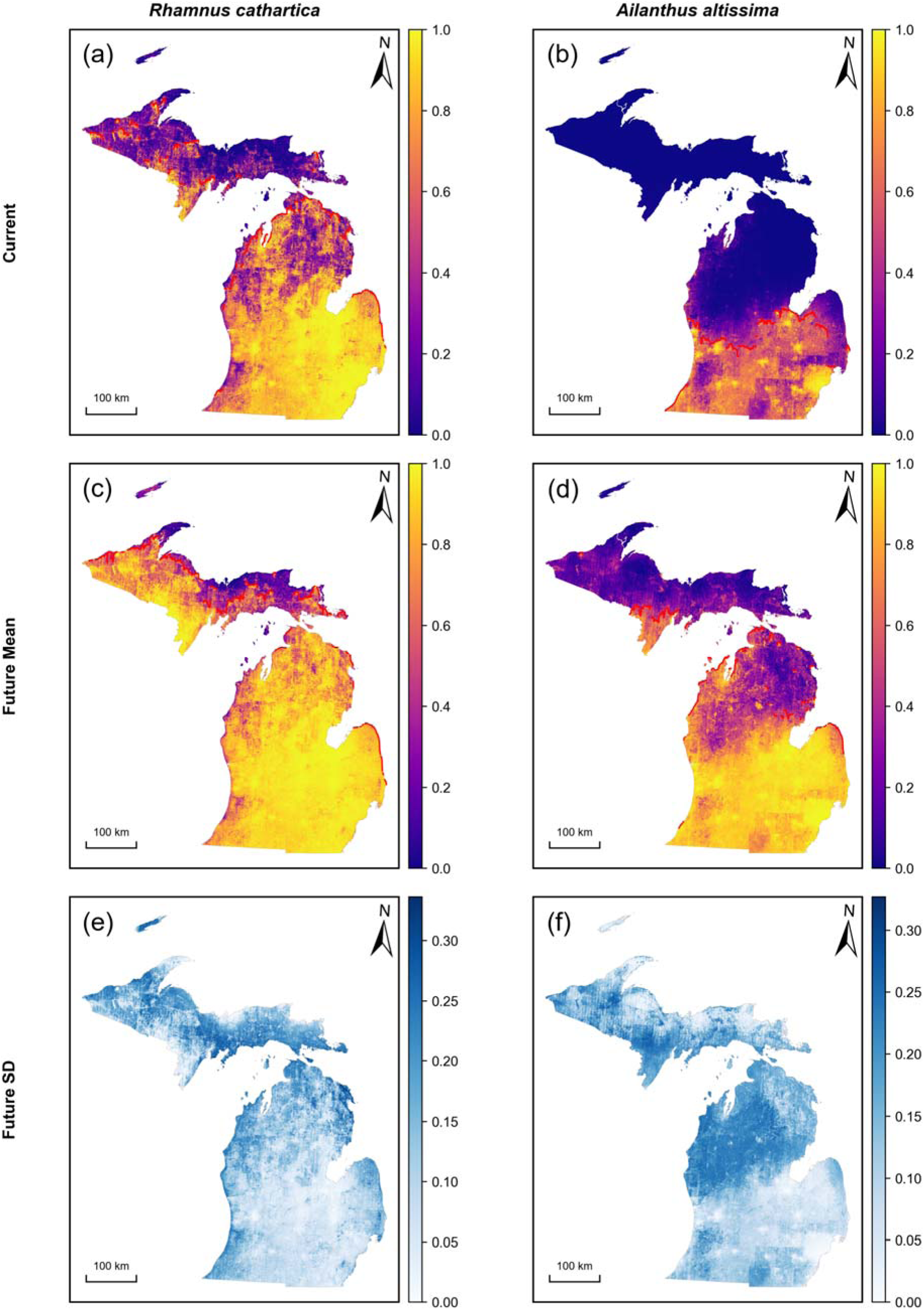
Maps of current, projected future habitat suitability, and projected future SD for two invasive species in Michigan. (a&b) Current probability of presence, (c&d) mean projected future probability of presence across climate scenarios, and (e&f) standard deviation of future predictions. Red lines represent the upper leading edge, defined as the northernmost pixel with predicted probability p ≥ 0.5 in each column.

**Table 2.**
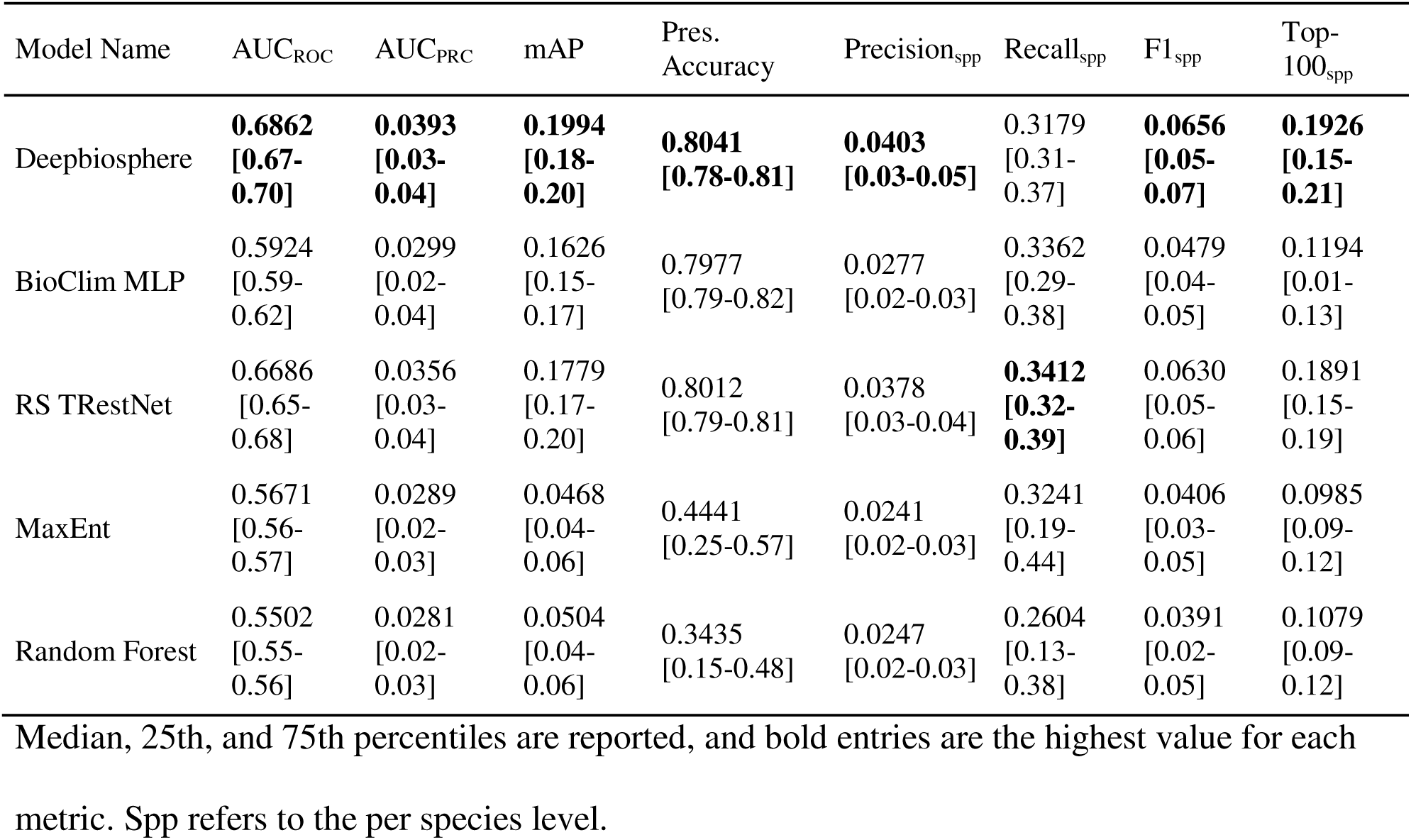
Comparison of accuracy metrics across 7 spatial cross-validation bands.

### 3.2 Current distribution maps

Current distributions for two invasive species were generated using the Deepbiosphere model, which outperformed other models (**Table 3**). For two invasive species, *R. cathartica* and *A. altissima*, Deepbiosphere improved modeling performance by an average of 56.41% and 74.99% in AUC-ROC, respectively. Compared with the 10.98% improvement across all species, these large increases suggested Deepbiosphere’s enhanced predictive capability for invasive species. Current habitat suitability maps showed clear differences between the two focal invasive species across Michigan (**Figure 3a&b**). *R. cathatica* exhibited a broadly high suitability pattern across most of the Lower Peninsula, with particularly high probability of presence in the southern and southeastern parts of Michigan. Its suitability gradually declined as it moved to the northern parts of Michigan in the Upper Peninsula, although some patches still had moderate to high suitability in the Upper Peninsula (**Figure 3a**). In contrast, *A. altissima* showed a more spatially restricted current distribution. High suitability was concentrated mainly in southern Michigan, especially in the southern Lower Peninsula, while most of the northern Lower Peninsula and Upper Peninsula showed extremely low suitability. The presence of *A. altissima* is hardly detected in the Upper Peninsula in the current day (**Figure 3b**). Overall, the current maps suggested that *R. cathatica* has a broader potential current distribution in Michigan, whereas the potential current distribution of *A. altissima* is more limited to the southern portion of Michigan.

**Table 3.** AUC-ROC values for *R. cathartica* and *A. altissima* across 5 models and 8 splits.

| Band | Deepbiosphere | BioClim MLP | RS TRestNet | MaxEnt | RF |
| --- | --- | --- | --- | --- | --- |
| *Rhamnus cathartica* |  |  |  |  |  |
| Uniform | **0.8478** | 0.7885 | 0.7583 | 0.7112 | 0.8215 |
| 1 | 0.7192 | 0.7185 | 0.6168 | **0.7837** | 0.6426 |
| 2 | 0.6726 | 0.7249 | 0.5924 | 0.7318 | **0.7327** |
| 3 | 0.8679 | **0.8789** | 0.7652 | 0.7761 | 0.7773 |
| 4 | 0.6596 | 0.5719 | **0.6769** | 0.6694 | 0.5236 |
| 5 | **0.8767** | 0.7453 | 0.7779 | 0.5091 | 0.393 |
| 6 | **0.8673** | 0.7873 | 0.852 | 0.4625 | 0.7278 |
| 7 | NA | NA | NA | NA | NA |
| *Ailanthus altissima* |  |  |  |  |  |
| Uniform | **0.7662** | 0.6575 | 0.7408 | 0.7135 | 0.6702 |
| 1 | **0.7177** | 0.6179 | 0.7092 | 0.6358 | 0.6693 |
| 2 | **0.7481** | 0.734 | 0.6568 | 0.6313 | 0.6709 |
| 3 | **0.9418** | 0.9117 | 0.8883 | 0.8073 | 0.9373 |
| 4 | 0.7888 | 0.8154 | 0.7956 | **0.82** | 0.8053 |
| 5 | **0.9998** | 0.8832 | 0.9777 | 0.3164 | 0.5933 |
| 6 | NA | NA | NA | NA | NA |
| 7 | NA | NA | NA | NA | NA |

Bold entries are the highest value for each split across models. “NA” indicates no test
observations were available in that band.

### 3.3 Future suitability prediction

Under future climate scenarios, the ensemble mean habitat suitability increased for both invasive species, although the magnitude and geographical distribution of change were different between them (**Figure 3c& d; Figure 4**). For *R. cathartica*, future mean habitat suitability remained high across the majority of the state and expanded northward further into the Upper Peninsula, indicating a broadly widespread invasion risk in the future. Compared with these areas with increasing risk, some parts of the eastern Upper Peninsula remained relatively low in habitat suitability. For *A. altissima*, the increase in predicted suitability was more pronounced than its current distribution. Although the current distribution was only concentrated in the southern Lower Peninsula, future mean habitat suitability showed a substantially broader range across much of the entire Lower Peninsula and expanded into the southern Upper Peninsula, occupying a relatively small area.

The change map further clarified these future shifts by showing the spatial distribution of increasing, stable, and decreasing suitability (**Figure 4c&d**). For *R. cathartica*, the stable areas were located in the southern and southeastern portions of the Lower Peninsula, exactly where high suitability areas in current distribution were located, which means high suitability remained in this region. Increasing suitability dominated the majority of the Upper Peninsula and much of the northwestern and northern parts of the Lower Peninsula. Decreasing areas were extremely limited on the very northern tip of the Upper Peninsula. Consistent with these spatial patterns, the area-based summary indicated that 55.36% of the state was classified as increasing suitability, 44.57% as stable, and only 0.007% as decreasing **(Table 4)**. In contrast, *A. altissima* exhibited a more widespread increase signal across most of Michigan, with much fewer stable areas than that of *R. carthartica* and no decreasing area. Only parts of the northern Upper Peninsula remained stable with low suitability. The invasion risk of *A. altissima* became more spatially extensive but still concentrated in the Lower Peninsula, where future mean suitability was relatively high. The area-based summary further confirmed this stronger expansion pattern: 86.39% of the state was classified as increasing suitability and 13.61% as stable, with no decreasing areas identified **(Table 4)**. Overall, these spatial change patterns indicate that both species are likely to expand northward under future climate conditions, but the increasing invasion risk is especially strong for *A. altissima* across the state.

**Figure 4.**
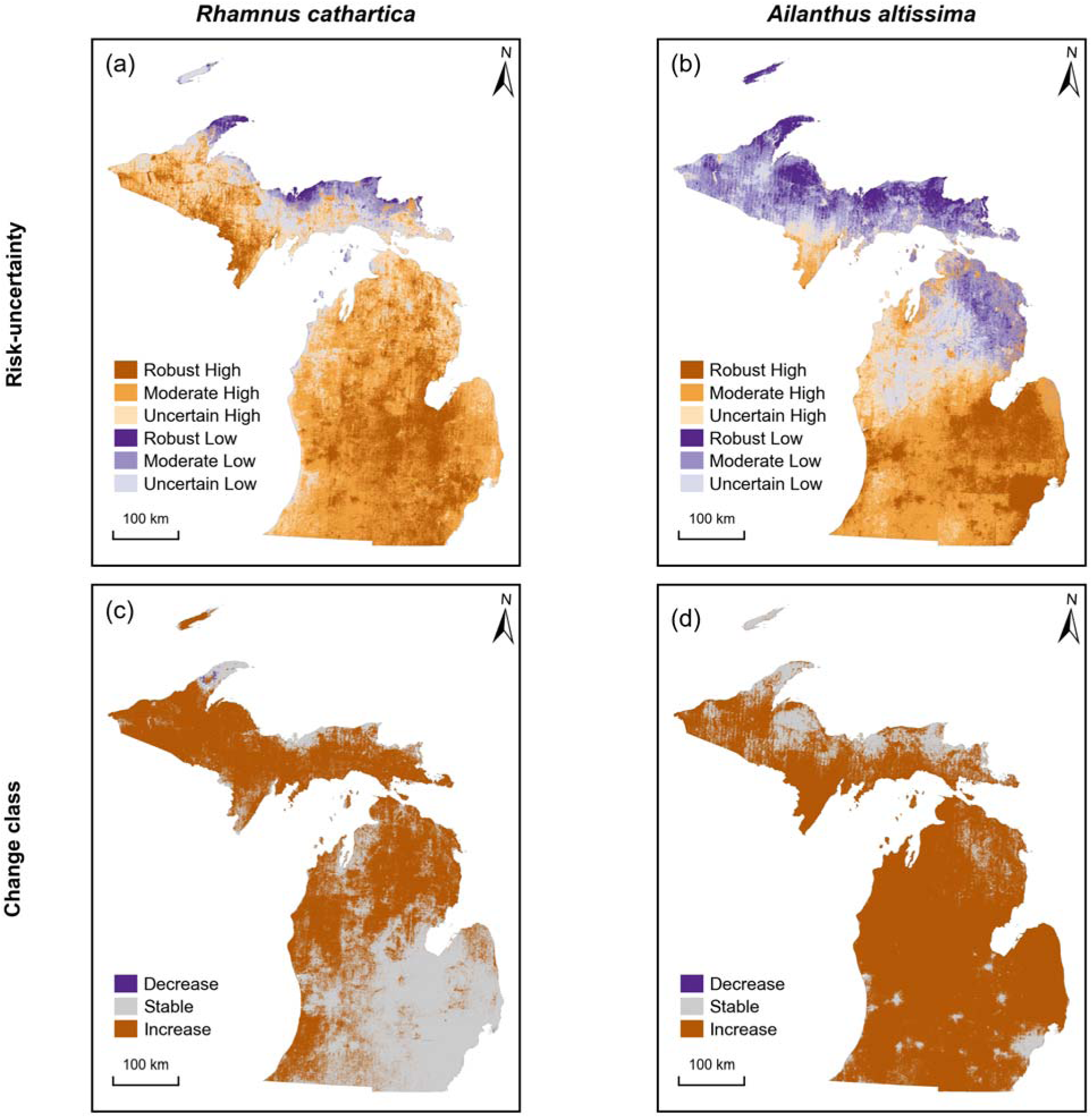
Maps of classification for risk-uncertainty and invasion risk change for two invasive species in Michigan. (a&b) Six-category risk–uncertainty map classifying areas into robust, moderate, and uncertain high-risk or low-risk zones; (c&d) invasion risk change map classifying probability into decrease, stable, or increase based on the difference between the current and future mean habitat suitability.

**Table 4.** Area-based summary of predicted habitat suitability change, dominant uncertainty source and risk-uncertainty classification for two invasive species in Michigan.

| Species | Change category | Proportion of total area (%) | Uncertainty source | Proportion of total area (%) | Risk-uncertainty Classification | Proportion of total area (%) |
| --- | --- | --- | --- | --- | --- | --- |
| *R. cathartica* | Decrease | 0.007 | GCM | 73.90 | Robust high | 23.7 |
|  |  |  |  |  | Moderate high | 46.5 |
|  | Stable | 44.57 | SSP | 2.21 | Uncertain high | 16.4 |
|  |  |  |  |  | Robust low | 1.3 |
|  | Increase | 55.36 | Residual | 23.89 | Moderate low | 3.5 |
|  |  |  |  |  | Uncertain high | 8.6 |
| *A. altissima* | Decrease | 0.00 | GCM | 95.9 | Robust high | 17.2 |
|  |  |  |  |  | Moderate high | 30.2 |
|  | Stable | 13.61 | SSP | 1.51 | Uncertain high | 10.2 |
|  |  |  |  |  | Robust low | 7.8 |
|  | Increase | 86.39 | Residual | 2.59 | Moderate low | 19.8 |
|  |  |  |  |  | Uncertain high | 14.8 |

### 3.4 Climate scenario uncertainty

Although both invasive species showed overall increases in future habitat suitability, the degree and source of uncertainty varied across Michigan. The standard deviation maps revealed clear spatial heterogeneity in uncertainty for both species, with relatively higher uncertainty concentrated in northern Michigan and transition zones between Upper and Lower Peninsula, whereas many areas in southern Michigan showed lower uncertainty under future climate scenarios (**Figure 3e&f**).

The risk-uncertainty maps further clarified how future invasion risk and uncertainty overlapped spatially (**Figure 4a& b; Table 4**). For *R. cathartica*, high-risk areas dominated most of the state, including 23.7% robust high risk, 46.5% moderate high risk, and 16.4% uncertain high risk, totaling 86.6%. Low-risk areas were limited for *R. cathartica*, with only 1.3% robust low risk, 3.5 moderate low risk, and 8.6% low risk. In contrast, low-risk areas were more extensive for *A. altissima*, 42.4% in total, including 7.8% robust low risk, 19.8% moderate low risk, and 14.8% uncertain low risk. However, the invasion risk of *A. altissima* still remained high for the rest 57.6% of the state, which included 17.2% robust high risk, 30.2% moderate high risk, and 10.2% uncertain high risk. For both species, robust and moderately high-risk areas were concentrated mainly in southern Michigan, whereas uncertain or low risk areas were more common in northern and transitional regions.

### 3.5 Variance decomposition of uncertainty

To partition the sources of uncertainty, the variance decomposition was conducted after fitting the additive model at the pixel level. The results showed that the sources of uncertainty were not evenly distributed across space (**Figure 5**). For both species, GCM was the dominant source of uncertainty across most of Michigan. For *R. cathartica*, GCM dominated 73.90% of the total area, followed by residual variation (23.89%), whereas SSP contributed only 2.21% (Table 3). For *A. altissima*, GCM dominance was even stronger, accounting for 95.90% of the total area, while SSP and residual components contributed only 1.51% and 2.59%, respectively (**Table 4**). These results show that uncertainty in future invasion risk projections was driven primarily by differences among climate models rather than by differences among SSP scenarios.

**Figure 5.**
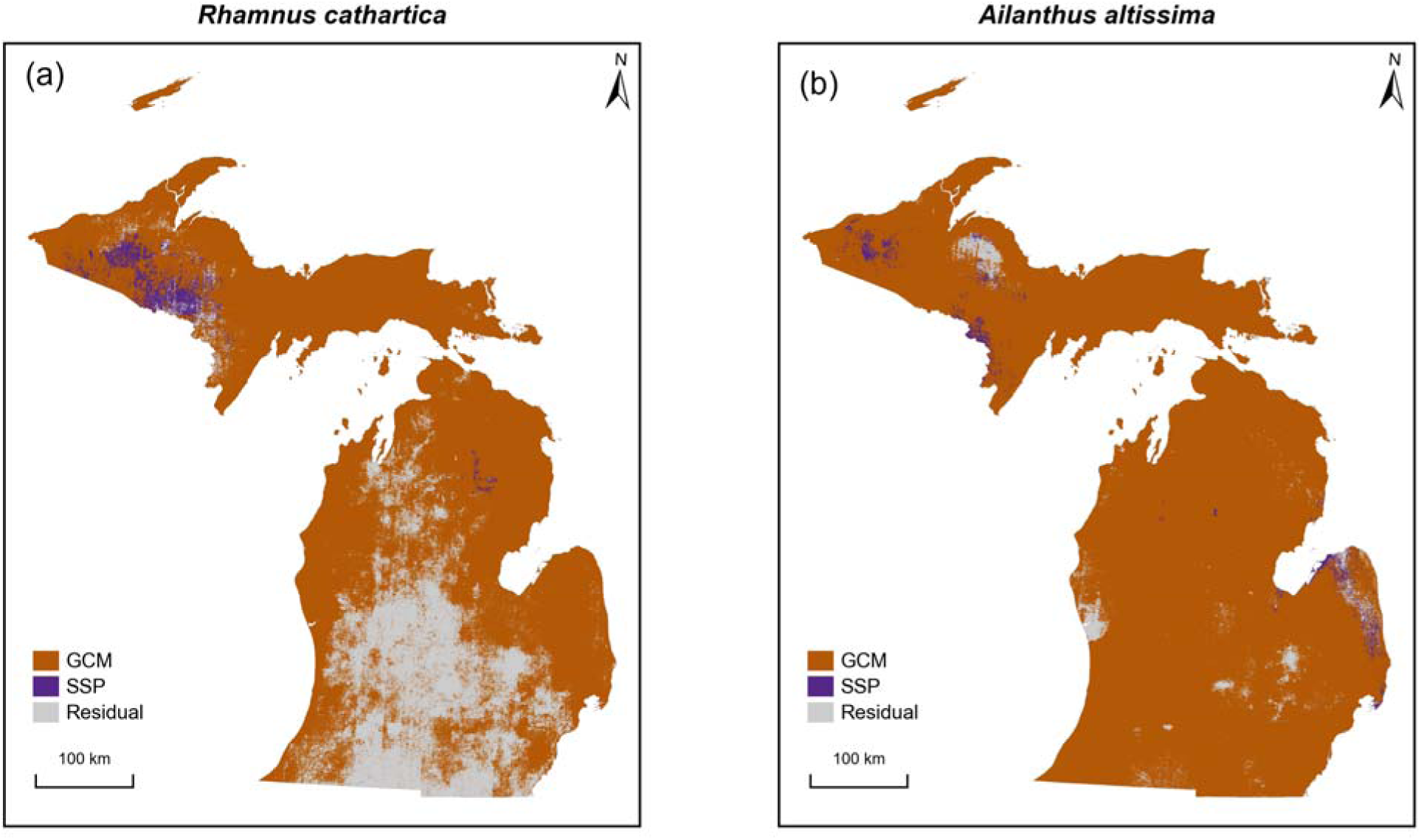
Dominant source of climate uncertainty for (a) *R. cathartica* and (b) *A. altissima* at each pixel in Michigan. Uncertainty sources are GCMs, SSPs, and Residual.

## 4 Discussion

### 4.1 Performance of Deepbiosphere at regional scales

The results of model comparison suggest that Deepbiosphere can be successfully transferred beyond its original region of California and can be effective for multispecies distribution modeling for more than 1000 plant species in Michigan. Deepbiosphere outcompeted other models for most metrics, either on the uniform split or spatial cross-validation splits.

Specifically, Deepbiosphere also achieved the strongest performance for two target invasive species. Another notable result was the large improvement in ranking metrics, especially mAP, which evaluated multi-label classification tasks where many species are predicted simultaneously. The higher performance of Deepbiosphere on these metrics suggests that the model was able to capture multispecies structure that is difficult to recover with single-species approaches, including local co-occurrence patterns and taxonomic relationships. This advantage was also supported by recent studies. Sharma et al. (2025) argued that multispecies and joint-species approaches can improve predictions by “borrowing strength” across species through shared information, rather than treating each species as an isolated prediction problem. Brun et al. (2024) found that multispecies deep learning models trained on citizen science data produced more informative plant community models. An invasive species study similarly showed that integrating surrogate co-occurring species within a joint species distribution modeling framework can improve predictions of range expansion (Briscoe Runquist et al., 2021), highlighting the value of co-occurrence information for forecasting invasive plant distributions.

In addition, the comparable performance of Deepbiosphere and TresNet on spatial cross-validation splits indicated that high-resolution NAIP imagery alone captures a large amount of predictive signal relevant to plant distributions in Michigan. For example, fine-scale land cover, vegetation texture, edge habitats, and human disturbance are all likely present in the imagery and may explain why the model with only remote sensing as input performed unexpectedly well.

Randin et al. (2020) emphasized that remote sensing products can provide continuous spatiotemporal information on key processes that drive species distributions, including land-cover dynamics, climate variability, and disturbance, thereby improving both the predictive and explanatory power of SDMs. In the context of invasive species, the ability to detect land-cover changes and disturbances is important because these conditions are where invasions are more likely to occur. In short, Deepbiosphere was still the most optimal model, suggesting that combining imagery with climate data and multispecies learning with multi-labels can improve plant distribution modeling. These architectural features are likely to be especially helpful in Michigan, where many species observations are relatively sparse compared with California (Gillespie et al., 2024).

### 4.2 Current and future invasion patterns

The current and future suitability maps suggest that the two target invasive species represent different invasion dynamics in Michigan. *R. cathartica* already showed a broad geographical distribution across much of the Lower Peninsula, indicating that it is likely a well-established invader under current environmental conditions. This is consistent with common buckthorn’s botanical description as a broadly adaptable invasive shrub capable of occupying both disturbed and relatively undisturbed habitats, including roadsides, old fields, and a variety of woodlands (MDAR, 2025). In contrast, *A. altissima* was more restricted under current conditions but showed a much stronger future expansion signal, with suitability projected to increase across most of Michigan. *A. altissima* is a thermophilic invasive species that benefits from warmer conditions and readily expands in disturbed and urban environments. A recent study similarly projected future increases in suitability and expansion risk for *A. altissima* under climate change, suggesting that warming may release climatic constraints on its further spread (Ding et al., 2022).

A common pattern across the two species was a tendency toward northward expansion with climate change, which aligned with many previous studies’ findings (Bradley et al., 2010; Bradley et al., 2024; Chen et al., 2011; Lenoir & Svenning, 2014). These studies found that global warming can increase habitat suitability and drive poleward or upslope range expansion, and invasive species are often favored by warming over native species, although the magnitude of the response may be species-specific. However, it should be noted that the potential of northward range shift does not necessarily mean that suitable habitat in the future will face high invasion risk, which can be demonstrated by *A. altissima* showing increasing suitability throughout Michigan but future high-risk areas will still remain more concentrated in the Lower Peninsula.

Another pattern worth noting was that many current and future hotspots facing high invasion risk were particularly located around urbanized areas and disturbed sites, such as the largest metropolitan area in the LP-Detroit, the largest city in the UP-Marquette, and a disturbed site, White Pine Copper Mine. High-risk pixels are concentrated in the city of Marquette and in the tailing basins of the White Pine Mine (**Figure 6**). This pattern is ecologically plausible because both target species are strongly associated with disturbed environments, with *A. altissima* especially characteristic of urban and industrial environments. Urbanized regions may provide conditions favorable for invasion, including higher propagule pressure, dense road networks that act as dispersal corridors, frequent disturbance, and local warmer environments caused by the urban heat island effect (Gaetner et al., 2017). This finding again supported that remote sensing may be able to represent these fine-scale anthropogenic features more directly than coarse climate data.

**Figure 6.**
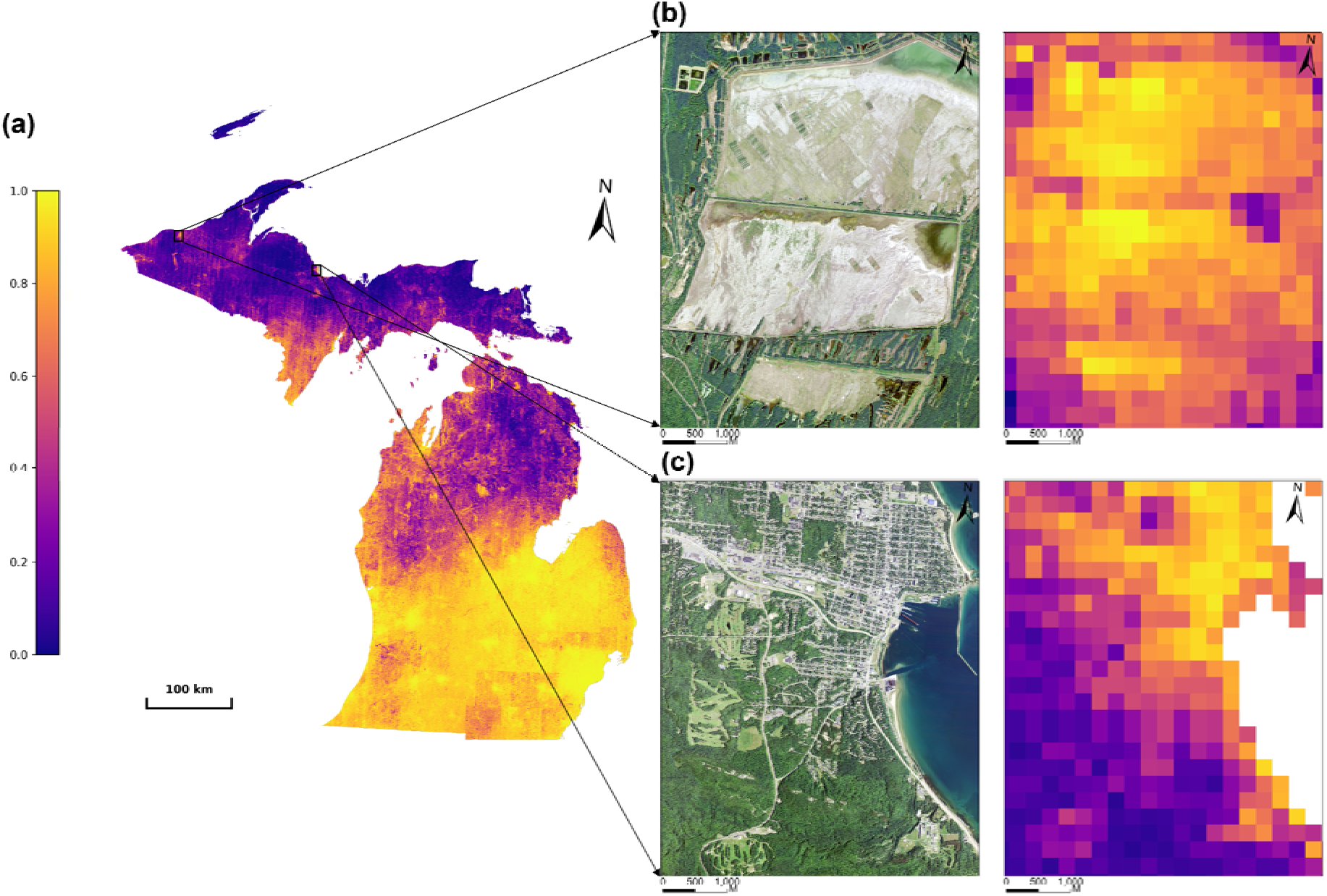
Two local zoom-in examples of future predictions for *A. altissima*. (a) Future mean habitat suitability of *A. altissima* in Michigan, (b) NAIP imagery and prediction at White Pine Mine, (c) NAIP imagery and prediction in Marquette.

### 4.3 Climate uncertainty and management implications

Our uncertainty analysis indicates that the climate scenarios influenced the predictions and their uncertainty in a spatially heterogeneous way, with different levels of uncertainty across Michigan for two species. Thomas et al. (2024) also conducted an invasive species case study to investigate the uncertainty of SDM algorithms and GCMs, but SSPs were not included. However, our variance decomposition results took SSPs into account and demonstrated that uncertainty from GCMs explained a larger proportion of variation than SSPs, becoming the dominant source of uncertainty for most areas in Michigan by the 2050s. This suggests that the choice of GCMs was a major driver of spatial disagreement in predicted invasion risk. Accordingly, future invasive species assessments in Michigan should prioritize the inclusion of multiple GCMs or select a certain GCM cautiously with sufficient justification in order to better communicate uncertainty.

From a management perspective, the risk–uncertainty framework provides guidance to inform invasive species management and conservation planning. Given that managing invasive species under climate change is challenging, we here propose several suggestions (**Table 5**).

**Table 5.** Conceptual framework for risk-uncertainty invasive species management.

|  |  | Risk |  |
| --- | --- | --- | --- |
|  |  | High | Low |
| Uncertainty | Robust | Immediate control<br>Large-scale removal<br>Habitat restoration<br>Long-term monitoring | Maintaining the native ecosystem integrity<br>Preventing new introduction |
|  | Moderate | Proactive containment<br>Preventing further spread and establishment | Low priority for direct intervention<br>Less frequent monitoring |
|  | Uncertain | Monitoring priorities<br>Early detection and rapid response (EDRR) | Field surveys to collect more data |

Robust high-risk areas may need immediate control, large-scale removal, and habitat restoration, coupled with long-term monitoring. Moderate high-risk areas may also warrant proactive containment, such as preventing further spread and establishment. Uncertain high-risk areas may be more appropriately treated as monitoring priorities, where early detection and rapid response are demanded. Robust low-risk areas should focus on maintaining the native ecosystem integrity and preventing new introductions. Moderate low-risk areas may be relatively low priority for direct intervention, where less frequent monitoring may be sufficient. Uncertain low-risk areas should not be considered as secure, because low-confidence areas are usually located in the Upper Peninsula, where data availability is limited, so field surveys may be needed to decrease uncertainty.

Compared with a GIS-based multicriteria decision analysis (MCDA) framework developed by Cohen et al. (2024), our risk-uncertainty framework serves a different but complementary purpose. The MCDA framework is a management-prioritization tool that ranks sites for invasive plant treatment by integrating documented invasive species occurrence with expert-weighted scores of integrity, biodiversity, resilience, and ecosystem services in Michigan. Thus, while the MCDA framework is designed to answer where management should be prioritized under current invasion, our risk-uncertainty framework aims to answer where invasion risk is predicted to increase and how confident those predictions are across future climate scenarios. Methodologically, these two frameworks differ in their construction. The variables used in MCDA are rule-based and expert-weighted, while our risk-uncertainty framework is heavily data-driven with empirical and statistical evidence. However, taken together, these two approaches can be highly complementary.

### 4.4 Limitations and future research

Despite the outstanding performance of Deepbiosphere in species distribution prediction, there are some systematic limitations in training data, model building, and future prediction. First, species occurrence data were derived primarily from citizen science observations, which are known to contain sampling bias. Although several preprocessing steps were applied to reduce bias, uneven sampling effort, and the lack of absence data may still have influenced model fitting and spatial prediction. Besides, the data size in Michigan was relatively smaller than that in California, where there were over 650,000 observations (Gillespie et al., 2024), whereas Michigan had only about 250,000 observations. A larger training dataset may enable further improvement in model performance (Wisz et al., 2008). Second, in terms of model building, unlike some studies utilizing species co-occurrence indices or matrices to infer ecological processes (Briscoe Runquist et al., 2021; Pollock et al., 2014), Deepbiosphere does not parameterize for such explicit biotic interaction terms, which may underestimate synergistic or allelopathic interactions among invasive and pioneer species. Third, our future predictions considered only the impact of climate change on habitat suitability and could not account for potential land use changes, which might be important for species distributions (Luo et al., 2025).

To overcome these limitations, future studies could focus on field surveys to collect absence data and independent ground truth data, especially in the data-sparse Upper Peninsula. Given that it is impossible to obtain remote sensing imagery for the future, future studies could also utilize land use projections as an extra predictor to substitute the lack of remote sensing data when predicting future species distributions (Frans & Liu, 2024; Luo et al., 2025; Newbold, 2018; Sinclair et al., 2010). In addition, future work could also improve the interpretability of Deepbiosphere predictions by applying Local Interpretable Model-agnostic Explanations (LIME) to examine which predictors drive local invasion-risk predictions. This may help explain more clearly whether predicted invasion risk is driven by anthropogenic disturbances from remote sensing or by climate change from environmental variables.

### 4.5 Conclusion

This study evaluated the applicability of a DL-based multispecies SDM framework, Deepbiosphere, for invasive plant species mapping in Michigan and examined how climate scenario uncertainty influences future invasion risk predictions. Deepbiosphere can be effectively applied to invasive plant species mapping in Michigan and can outperform baseline models across both interpolation and extrapolation settings. By integrating citizen science observations, high-resolution remote sensing imagery, and climate data, the model was able to accurately generate 256-meter resolution statewide predictions for *R. cathartica* and *A. altissima* and revealed clear differences in their current and future invasion patterns. Under future climate scenarios, both species were projected to expand northward, and future invasion risk would increase, accompanied by spatial heterogeneity in uncertainty. Additionally, GCMs were the dominant source of uncertainty across Michigan. Our risk-uncertainty framework offers an actionable basis for monitoring and management. Overall, this study highlights the value of combining deep learning with spatially explicit uncertainty analysis to support more informed invasive species management under climate change.

## Author Contributions

Xu Qiang: Conceptualization; methodology, writing-original draft; Lauren Gillespie: Supervision, methodology, writing-review and editing; Xi Jia: Post-hoc data analysis; Dimitrios Gounaridis: Supervision; writing-review and editing; Kai Zhu: Conceptualization, supervision, writing-review and editing

## Acknowledgement

We thank all the experts and citizen scientists who contributed plant observations to the GBIF database. Kai Zhu was supported by NSF award 2306198. We thank the members of the Zhu Lab for their valuable comments and feedback on the manuscript.

