## Supplementary Materials for "Mapping distribution of invasive plant species and uncertainty using citizen science, remote sensing, and deep learning"

### Methodology

Detailed information of data processing and justification for why and how we did are discussed below. The workflow of generating the final joint dataset is illustrated in **Figure S1**.

#### Species occurrence data

We collected plant occurrence data across the Kingdom Plantae in Michigan from the Global Biodiversity Information Facility (GBIF.org, 2025), covering the period from 2015 to 2025. We filtered the records to include only those observed by humans with a coordinate uncertainty radius of less than or equal to 120 meters and no geospatially flagged issues. In total, 305,343 observations across 2705 species,1009 genera, and 252 families were downloaded. We focused exclusively on vascular plants, so we filtered the dataset only to include the observations of Gnetopsida, Liliopsida, Lycopodiopsida, Magnoliopsida, Pinopsida, Polypodiopsida, and Ginkgoopsida. Then, following the preprocessing steps Gillespie et al. (2024), we conducted spatial thinning to reduce sampling bias by removing duplicate records of the same species within a 150 m radius, and eliminating species whose observations were confined to a single 256 m radius.

To create the joint occurrence dataset, we applied neighbor imputation, which added any additional species observations observed within an overlapping 256 m radius to a given observation. The joint occurrence dataset was motivated by the ecological theory supported the idea that biotic interactions strongly influence species’ distributions at many scales (Wisz et al., 2013) and that joint SDMs could predict species distributions more accurately than commonly used approaches (Brun et al., 2024; Clark et al., 2014). Finally, any species that contained fewer than 200 total observations were removed. This resulted in a dataset of 248498 total observations encompassing 1553 unique vascular plant species across 676 genera and 157 families.

#### Remote sensing Imagery

For remote sensing data, we used imagery from the National Agricultural Imagery Program (NAIP) acquired in 2012 for the entire state of Michigan (USGS Earth Resources Observation and Science Center, 2018). NAIP imagery provides 1-meter spatial resolution with as much as 10 percent of cloud cover in each tile, enabling detection of fine-scale landscape features and individual plant community structures that are often obscured in medium-resolution satellite data like Landsat (30m) or Sentinel-2 (10m). In terms of spectral resolution, NAIP Imagery contains RGB and Near-infrared bands, which provide critical vegetation information for the model to learn to distinguish species.

As demonstrated by Hansen et al. (2013), major shifts in forest cover and vegetation structure are relatively slow in the absence of catastrophic disturbances, justifying the use of 2012 NAIP imagery as a stable structural proxy for the study period. Ecological lag effects suggest that current species distributions are often determined by past habitat environments (Svenning & Sandel, 2013). Therefore, 2012 imagery can capture landscape features that were utilized by plants observed between 2015 and 2025. Besides, NAIP transitioned to a finer spatial resolution standard of 0.6 meters after 2017 (USGS EROS, 2018), meaning that storage and computing requirements would be prohibitively higher. Therefore, to balance storage constraints, computational feasibility, and model accuracy, we finally selected the remote sensing imagery from NAIP 2012.

The raw NAIP imagery was obtained as standard Digital Ortho Quarter Quads (DOQQs) covering the entire extent of Michigan, with each individual image tile covering a geographic area of a 3.75-minute longitude by 3.75-minute latitude quarter quadrangle (USGS EROS, 2018). However, to train the CNN model, the images were partitioned into 256m×256m tiles, where each pixel represented a 1m×1m. The NAIP quarter quads in Michigan are projected into two projection coordinate systems, depending on their location. Therefore, we transformed all tiles in UTM Zone 16N into UTM Zone 17N. Then, we linked each species observation by its geographic coordinates to a 256m×256m tile with the observation centered in the image.

#### Bioclimatic Variables

We downloaded the current 19 bioclimatic variables from WorldClim Version 2 at 30 arc-second resolution, which is approximately 1km spatial resolution (Fick & Hijmans, 2017). These variables are derived from monthly temperature and precipitation in order to generate more biologically meaningful variables, and they summarize annual trends, seasonality, and extreme or limiting climatic conditions. Bioclimatic variables from WorldClim were selected because they were biologically meaningful and were commonly used in SDM. All bioclimatic variables were standardized using z-score normalization to a mean of 0 and a standard deviation of 1 across Michigan.

For future projections, we obtained bioclimatic variables from multiple Global Climate Models (GCMs) under different Shared Socioeconomic Pathways (SSPs) for the mid-21st century (2041–2060). Future bioclimatic variables were also downloaded from WorldClim Version 2 at 30 arc-second resolution (Fick & Hijmans, 2017). To capture climate model uncertainty, we selected five GCMs that covered low to high climate sensitivity projections. The selected GCMs were MIROC6 (ECS ≈ 2.6°C), BCC-CSM2-MR (≈ 3.0°C), MRI-ESM2-0 (≈ 3.2°C), CMCC-ESM2 (≈ 3.6°C), and IPSL-CM6A-LR (≈ 4.5°C). To capture emissions pathway uncertainty, we selected three SSPs representing low, intermediate, and high greenhouse gas emission trajectories: SSP1-2.6, SSP2-4.5, and SSP5-8.5. Variation in change in predicted habitat suitability across these combinations was interpreted as climate scenario uncertainty.

#### Train/Test partitioning

Data partition, or data splitting, is a fundamental part of machine learning reproducibility. Normally, a dataset is divided into train, test, and validation, but in this study, we only split the dataset into train and test for two reasons. First, the validation dataset should have at least several thousand data points that are independent from the GBIF species occurrence data. Second, citizen science data are primarily collected in regions that are publicly or easily accessible, so the potential locations to collect validation data tend to be private, unpublic, or remote lands (Gillespie et al., 2024). Due to these two constraints, it would be difficult for us to build an independent validation set; Instead, we focused on splitting the train and test set.

As widely discussed, it is more likely that spatially close data points share similar attribute values because of the spatial dependency (Tobler, 1970; Hijmans, 2012). To address this problem, we used two types of train/test splitting: uniform split and spatial cross-validation splits. The uniform split was to randomly select a subset of observations uniformly across Michigan as the test set, while the remaining observations were used for training. To mitigate data leakage due to spatial autocorrelation, we put all overlapping observations in the test set, so that the images and observation data of the test set were excluded from the train set. Besides, as the diameter of a pixel in bioclimatic raster could be up to 1200m converted from arc-second, we removed the observations that were located within 1300 m of observations in the test set, thus avoiding the same input value of bioclimatic variables in both the train and test sets.

In the absence of an independent validation dataset, we further used spatial cross-validation to test model extrapolation ability. The spatial cross-validation split involved a 7-fold cross-validation. The Michigan study region was partitioned into several latitudinal bands (**Figure S2**). In each fold of the spatial cross-validation, one band was held out as the test region while the remaining bands were used for training. After several trials setting the bandwidth to 0.5, 0.7, 1, and 2 degrees, 1 degree was chosen because the ratio of each train/test set using 1 degree was the most reasonable. As discussed above, training observations within 1300 m of the test region were also removed to prevent data leakage. Due to differences in observation density across geographic bands, the number of training and testing samples varied among 7 cross-validation folds **(Table S1**).

#### Model training

We trained all deep learning models using a learning rate of 1×10⁻^5^, 12 epochs, and a batch size of 150. We explored multiple fixed learning rates, including 1×10⁻^6^, 1×10⁻^5^, and 1×10⁻^4^, and compared them with a cosine annealing learning rate schedule from 1×10⁻^6^ to 1×10^-2^ on the uniform split. Preliminary experiments indicated that a fixed learning rate of 1×10⁻^5^ provided the best training performance based on AUCROC and loss. Therefore, we adopted a fixed learning rate of 1×10⁻^5^ in the final model training. We used a recent loss function, sampling-aware binary cross-entropy (BCE), to measure the discrepancy between predicted probabilities and observed species occurrence labels. Accounting for sampling bias in citizen science data, sampling-aware BCE has been demonstrated to perform well when dealing with multi-label image classification tasks because it ensures that present and absent classes contribute equally to the total loss. (Gillespie et al., 2024). Model configuration and input data used to train Deepbiosphere and other baseline models are presented in **Table S2**.

#### Uncertainty analysis

Our study evaluated future invasion risk together with prediction uncertainty. The use of ensemble mean suitability reflects the fact that the future climate cannot be certainly known. Rather than relying on any single scenario, averaging future suitability across multiple climate scenarios provides a summary of central tendency, while preserving the opportunity to assess uncertainty among scenarios. This climate uncertainty analysis was justified by previous studies that uncertainty should be explicitly communicated and quantified in species distribution prediction under climate change (Beaumont et al., 2008; Gould et al., 2014). Buisson et al. (2010) showed that uncertainty in future predictions can be partitioned among various sources that might be spatially heterogeneous.

To specifically evaluate the effect of future climate scenarios on predicted species presence and associated uncertainty, we analyzed changes in predicted presence probability relative to the current baseline rather than raw future predictions. Raw future predictions reflect both current spatial patterns, including current landscape structure captured by remote sensing imagery, and changes induced by future climate conditions. By subtracting the current prediction from each future prediction, the resulting ΔP values represent the direction and magnitude of projected change. This approach therefore reduces the influence of baseline spatial patterns and allows the uncertainty analysis to focus more directly on scenario-driven variation associated with future climate scenarios.

An additive model was used to decompose variance because each GCM–SSP combination produced only one predicted value for each species at each pixel, which means there was no replicated predictions within the same GCM–SSP combination. As a result, there was no independent within-group variation that could be used to estimate a separate interaction term. Therefore, after accounting for the additive contributions of GCMs and SSPs, all remaining unexplained variation was assigned to the residual component.

### Post-hoc Covariate Shift Analysis

Covariate shift analysis was conducted to evaluate whether differences in model performance among spatial cross-validation bands were associated with differences in the environmental conditions represented in the training and testing sets. The cross-validation splits contained 7 latitudinal folds, which were naturally correlated with climate gradient, land use/land cover patterns and vegetation types in Michigan. As a result, cross-validation splits tested not only the model’s ability to generalize to unseen samples, but also its ability to extrapolate to environmental conditions that may be different from the training data. Therefore, we compared the statistical distribution of training and testing sets for each split. If the testing set has a distinct distribution from the training set, model predictions become more extrapolative and should be interpreted with greater caution. This is especially important for explaining the variation in spatial cross-validation performance, because reduced accuracy in a given band may indicate new environment and vegetation types or sample imbalance rather than model failure in extrapolation.

The covariate shift analysis showed that most spatial cross-validation bands had relatively comparable training and testing covariate distributions, whereas Band 7 showed a mismatch in statistical distribution between the training and testing data of both environmental data and remote sensing data (**Figure S2&S3**). Additionally, Band 7 had the most imbalanced number of training and testing samples and the fewest shared species between train and test sets (**Table S1**), which meant that there might be no testing sample for many species. As a result, it was reasonable that Band 7 showed the poorest performance and exhibited outlier behavior in the spatial cross-validation results. To avoid this single extreme band disproportionately influencing the overall model comparison, median performance across spatial cross-validation bands was reported instead of mean performance.

### Figures and Tables

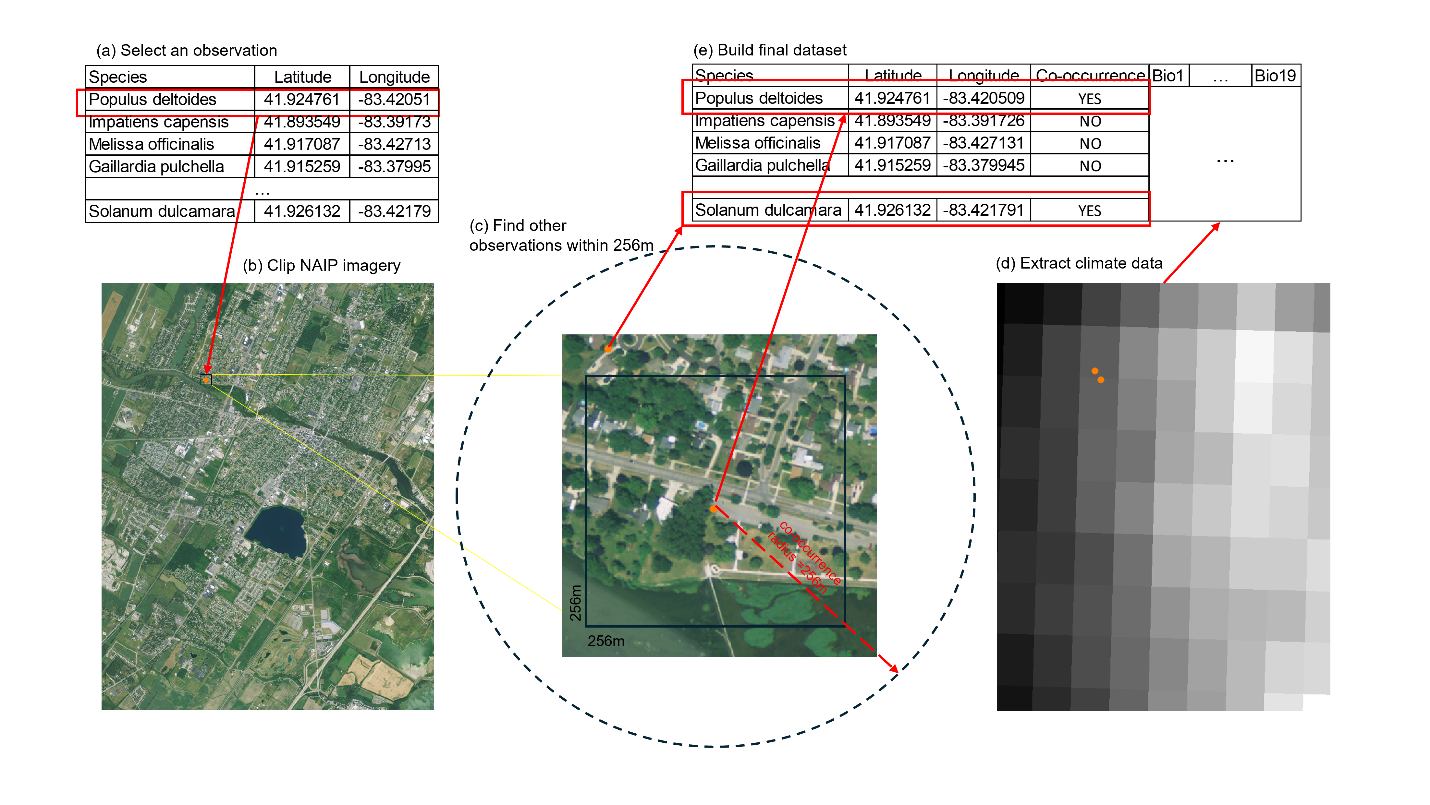

**Figure S1.** Illustration of building the final dataset. (a) Select a random observation record from GBIF. (b) Clip NAIP imagery with the observation centered at the clipped tile. (c) Find other species observations within 256m of the given observation to create a joint occurrence dataset. (d) extract bioclimatic variables for co-occurring observations. (e) Build final dataset.

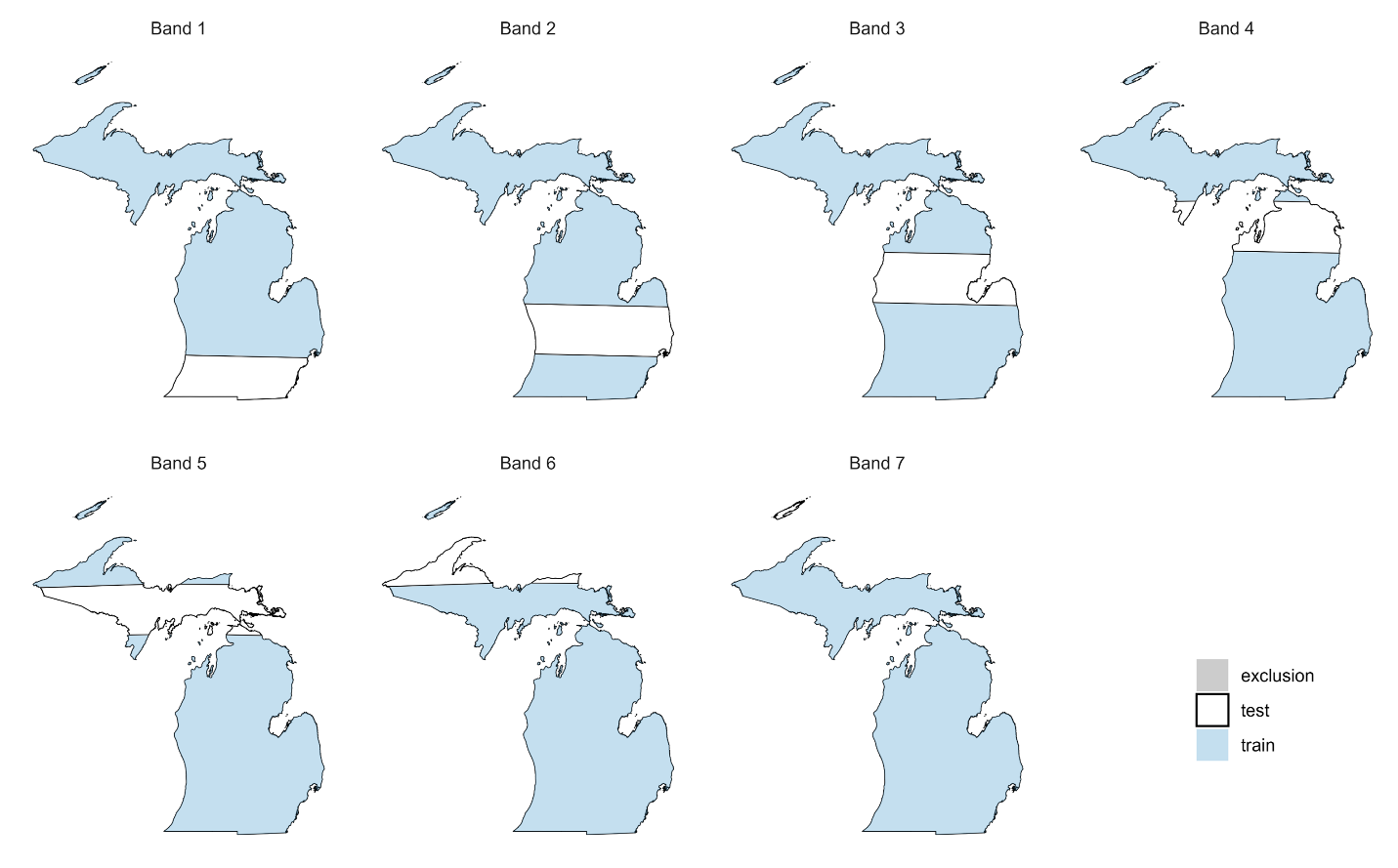

**Figure S2.** Visualization of 7-fold spatial cross-validation. White areas are the testing set, blue areas are the training set. Locations where observations are within 1.3km of the test region are removed from the training set and are colored in grey.

**Table S1.** Summary of training and testing partitions.

| Partition Type | Latitude Range | Train Count | Train % | Test Count | Test % | Total | Shared Species |
| --- | --- | --- | --- | --- | --- | --- | --- |
| Uniform | NA | 236461 | 95.16% | 12037 | 4.84% | 248498 | 1123 |
| Band 1 | 41.5-42.5 | 179169 | 72.10% | 68357 | 27.51% | 247526 | 1298 |
| Band 2 | 42.5-43.5 | 176386 | 70.98% | 70961 | 28.56% | 247347 | 1341 |
| Band 3 | 43.5-44.5 | 234496 | 94.37% | 13587 | 5.47% | 248083 | 1033 |
| Band 4 | 44.5-45.5 | 211293 | 85.03% | 36661 | 14.75% | 247954 | 1226 |
| Band 5 | 45.5-46.5 | 216184 | 87.00% | 30660 | 12.34% | 246844 | 1088 |
| Band 6 | 46.5-47.5 | 226306 | 91.07% | 21212 | 8.54% | 247518 | 983 |
| Band 7 | 47.5-48.5 | 247158 | 99.46% | 1340 | 0.54% | 248498 | 234 |

Latitudinal partition range, numbers and percentages of training and testing observations, and the number of species shared between training and testing sets are reported.

**Table S2.** Model configuration includes input data, hyperparameter and loss function used for training.

| Model Name | Input Data Type | Learning Rate | Batch Size | Epochs | | Loss Function |
| --- | --- | --- | --- | --- | --- | --- |
| DeepBiosphere | NAIP & Bioclim | 1e-5 (0.00001) | 150 | 12 | | Sampling-Aware BCE |
| Bioclim MLP | Bioclim Only | 1e-5 (0.00001) | 150 | 12 | | Sampling-Aware BCE |
| RS TResNet | NAIP Only | 1e-5 (0.00001) | 150 | 12 | | Sampling-Aware BCE |
| Random Forest | Bioclim Only | 1000 trees  No threshold | | | Default settings | |
| MaxEnt | Bioclim Only |  |  |  |  |  |

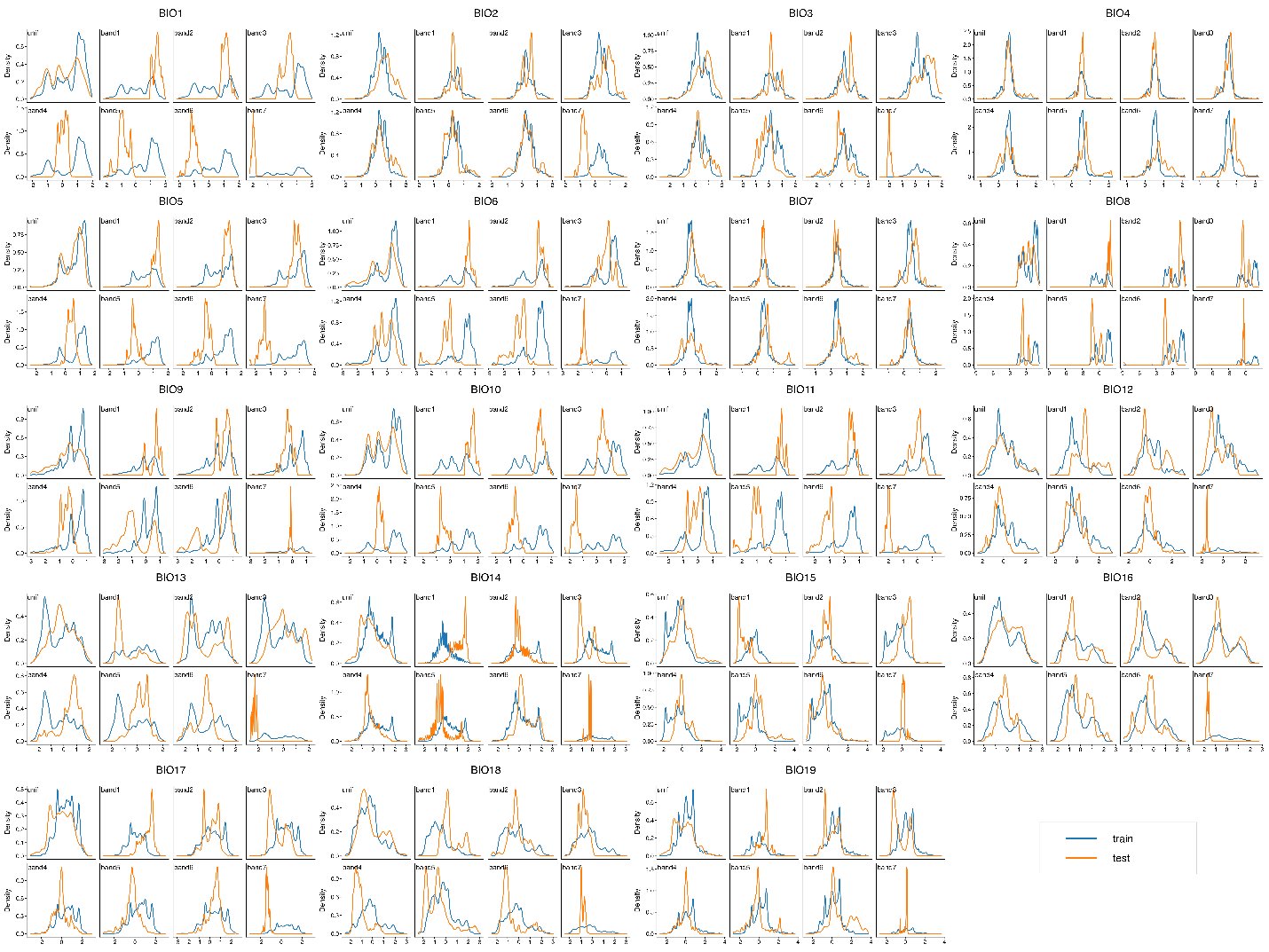

**Figure S3.** Statistical distribution of train/test data for 19 bioclimatic variables across 8 splits, showing distribution shift occurred with Band 7.

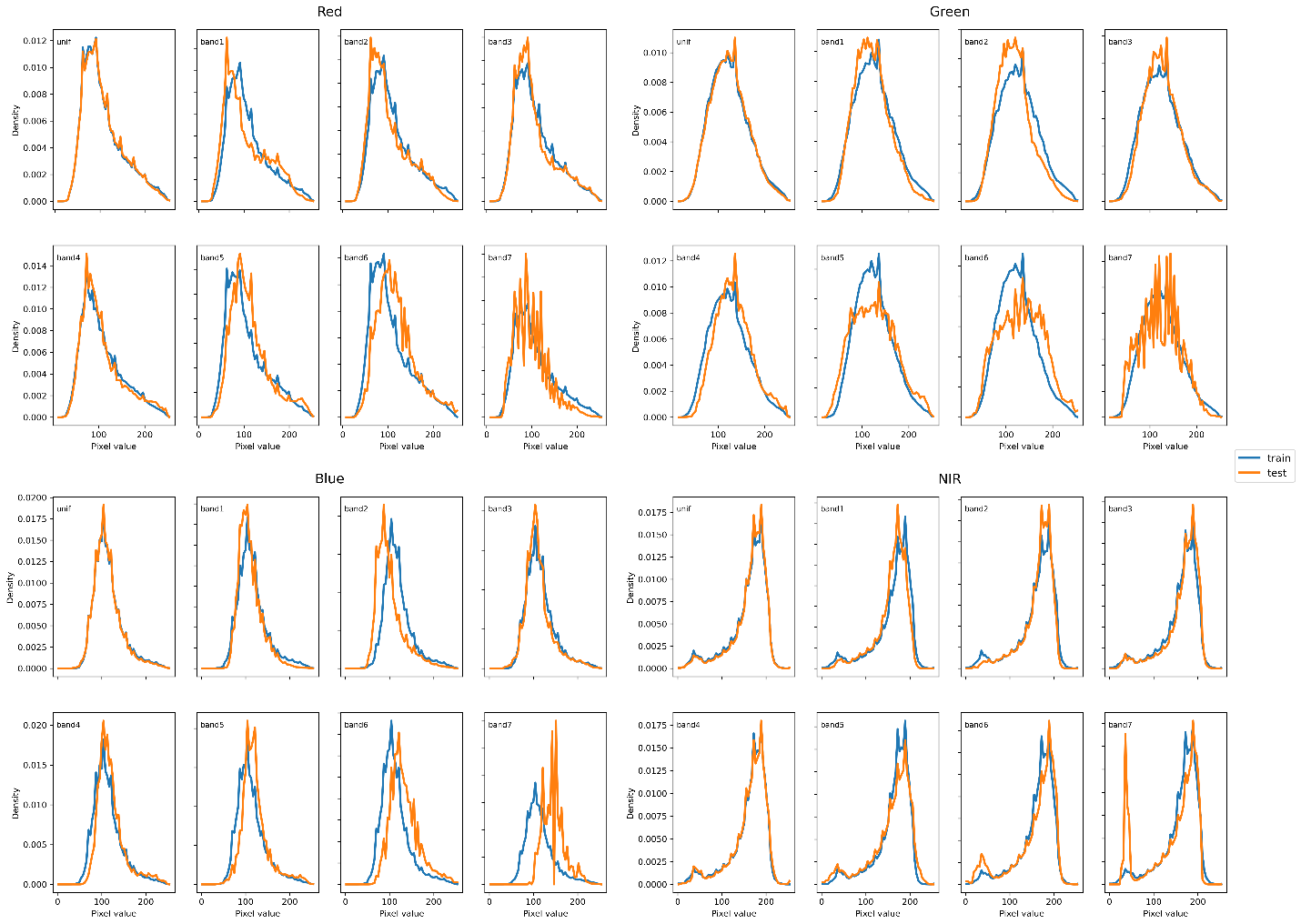

**Figure S4.** Statistical distribution of train/test data for Red, Green, Red, and NIR bands across 8 splits, showing distribution shift occurred with Band 7.
